# CA2 inhibition reduces the precision of hippocampal assembly reactivation

**DOI:** 10.1101/2020.11.26.400655

**Authors:** Hongshen He, Roman Boehringer, Arthur J.Y. Huang, Eric T.N. Overton, Denis Polygalov, Kazuo Okanoya, Thomas J. McHugh

## Abstract

The structured reactivation of hippocampal neuronal ensembles during fast synchronous oscillatory events termed sharp-wave ripples (SWRs) has been suggested to play a crucial role in the storage and use of memory. Activity in both the CA2 and CA3 subregions can proceed this population activity in CA1 and chronic inhibition of either region alters SWR oscillations. However, the precise contribution of CA2 to the oscillation, as well as to the reactivation of CA1 neurons within it, remains unclear. Here we employ chemogenetics to transiently silence CA2 pyramidal cells in mice and observe that while SWRs still occur, the reactivation of CA1 pyramidal cell ensembles within the events lose both temporal and informational precision. These observations suggest that CA2 activity contributes to the fidelity of experience-dependent hippocampal replay.

## Introduction

The differential anatomy, physiology and connectivity of neurons in the distinct subfields of the hippocampus allows these regions to play distinct and synergistic roles in the circuit’s mnemonic functions, both during memory formation and consolidation(Buzsaki, 2015; Goode et al., 2020; Hainmueller and Bartos, 2020; Jones and McHugh, 2011; Kay and Frank, 2019; Kesner and Rolls, 2015; Knierim and Neunuebel, 2015; Middleton and McHugh, 2019; Nakazawa et al., 2004; Oliva et al., 2016b). Of particular interest has been the observation that the assemblies of CA1 place cells active during exploration can be reactivated off-line during subsequent periods of rest(Foster and Wilson, 2006; O’Neill et al., 2006; van de Ven et al., 2016; Wilson and McNaughton, 1994). This experience-dependent replay of place cell sequences during sharp-wave ripples (SWRs) has been suggested to play a crucial role in memory consolidation(Carr et al., 2011; Karlsson and Frank, 2009; Squire et al., 2015) and disruption of these oscillations can impair learning(Ego-Stengel and Wilson, 2010; Girardeau et al., 2009; Jadhav et al., 2012). However, the dynamic contributions of the individual subregions of the circuit to these events has been difficult to disambiguate.

The canonical model of SWR generation has focused on CA3 to CA1 transmission as the main excitatory drive(Buzsaki, 2015; Csicsvari et al., 2000) and acute optogenetic silencing of CA3 input dramatically decreases the occurrence of SWRs, eliminating both the oscillation and the underlying sequential spiking(Davoudi and Foster, 2019). Chronic CA3 silencing, however, results in a more nuanced phenotype, with SWRs still present at normal levels, albeit at a significantly lower intrinsic frequency(Middleton and McHugh, 2016; Nakashiba et al., 2009). Recent work has suggested that this may be because CA2 input can also trigger SWRs(Oliva et al., 2016a) in CA1. However, chronic CA2 silencing also alters the structure, but not the occurrence of SWRs, leading a fraction of the events to become pathophysiologically large and fast(Boehringer et al., 2017), possibly related to the mutually inhibitory nature of the bidirectional feed-forward inhibitory connections between CA2 and CA3. Further, the preferential involvement of CA2 in social rather than spatial coding(Hitti and Siegelbaum, 2014) has led to the suggestion that CA2 and CA3-generated SWRs may make differential contributions to these two types of memory. Indeed, optogenetic disruption of CA2-generated SWRs has been shown to disrupt consolidation of social but not spatial experience(Oliva et al., 2020).

Despite these advances, inconsistencies and questions remain. First, the optogenetic and electrical stimulation protocols used to disrupt SWRs abolish both the oscillation and the underlying CA1 spiking activity. This makes it impossible to distinguish the contribution of CA2 or CA3 input to the generation of the population event from their role in the precise temporal coordination of the structured spiking of the reactivated ensemble. Next, although SWRs can be generated by activity in either CA2 or CA3, the vast majority of events arise from coordinated activity in these adjacent and highly connected regions(Oliva et al., 2016a), thus chronic silencing approaches makes interpretation of the changes more challenging. Hence, here we employ acute DREADD-mediated silencing of CA2 pyramidal cell activity to ask if we can differentiate the role of CA2 in ripple generation from its possible distinct contribution to influencing which neurons participate in these reactivations, as well as to the events temporal precision.

## Results

To investigate the influence of CA2 transmission on both the generation of sharp-wave ripples (SWRs) and the underlying neuronal assembly activation a multi-channel drive array was implanted in CA2-Cre mice(Boehringer et al., 2017) bilaterally injected with Cre-dependent AAV expressing either the inhibitory DREADD receptor(Armbruster et al., 2007), hM4Di, and mCherry (DREADD) or mCherry alone (Ctl) as a control, with tetrodes targeting all CA fields of the hippocampus (**Fig. 1a; Extended Data Fig. 1a, b**). Recording started with a baseline rest session (Pre-CNO) followed by exploration of a linear track (LT1), after which Clozapine-N-oxidase (CNO), the ligand for the hM4Di receptor, was injected systemically (CNO session). Mice were then exposed to second linear track (LT2), followed by a final rest session (Post-CNO) (**Fig. 1b**). This protocol allowed us to compare neural activity before and after CA2 inhibition to address how CA2 impacts the hippocampal network’s ability to generate SWRs and how it influences their experience-dependent spike content.

**Figure 1.**
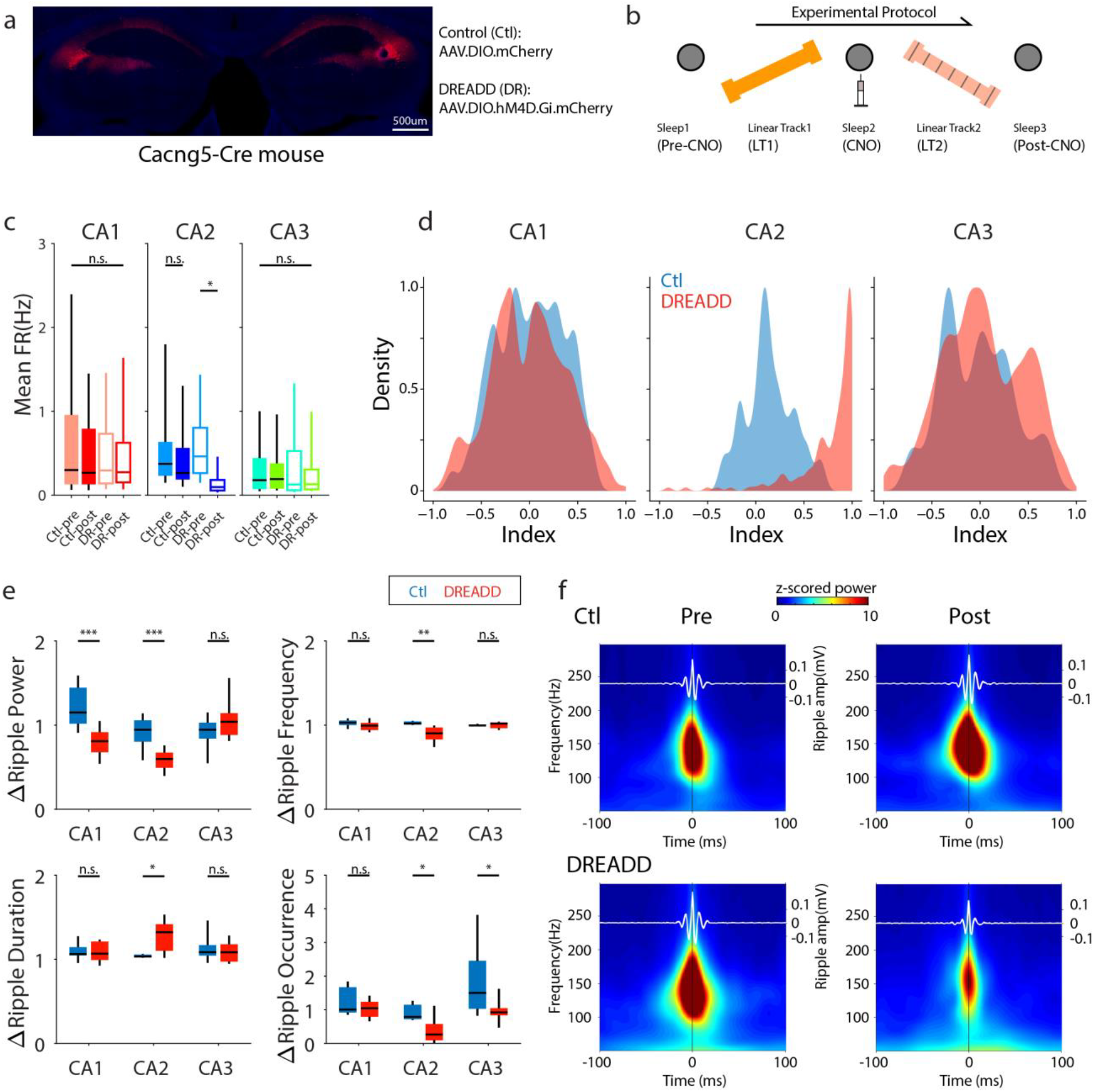
Acute silencing of CA2 pyramidal cells decreases SWR power. **a.** CA2-specific AAV-mediated mCherry expression in the Cacng5-Cre mouse. The lesion on the right side represents a CA2 recording site. **b.** Experimental design for assembly and replay analyses. **c.** Mean firing rate of pyramidal cells in the Pre- and Post-CNO session for control (filled; N=6 mice, 9 sessions) and DREADD (open; N=7 mice, 19 sessions) mice. A significant main effect of the session (*F*(1,234)=4.094, *P*=0.044) was found in CA2 only and post-hoc comparisons revealed a significant decrease in DREADD (pre>post, *P*=0.033) but not in control mice (pre vs. post, *P*=0.368). Two-way ANOVA with Bonferroni’s multiple-comparisons (see **Extended Data Table 1** for detailed statistics). **d.** Population density plot of the firing rate index, calculated as (FRPre-FRPost)/(FRPre+FRPost). A value of 1 represents a complete shutdown of firing. A significant difference was found in CA2 (*P*=1.06×10^−20^) but not in CA1 (*P*=0.66) or CA3 (*P*=0.65). Two-sample Kolmogorov-Smirnov test. **e.** Normalized changes in ripple properties between the Pre-CNO and Post-CNO session. Ripple power significantly decrease in CA1 (Ctl>DR: *P*=1.966×10^−11^, *Z*=6.708) and CA2 (Ctl>DR: *P*=0.011, *Z*=2.528), but not in CA3 (*P*=0.123, *Z*= −1.528) of the DREADD mice; ripple frequency showed a significant decrease in CA2 (Ctl>DR: *P*=0.009, *Z*=2.601), but not in CA1 (*P*= 0.070, *Z*=1.809) or CA3 (*P*=0.479, *Z*= −0.709); Ripple duration showed a significant increase in CA2 (Ctl<DR: *P*=0.040, *Z*= −2.054), but not in CA1 (*P*=0.760, *Z*=0.305) or CA3 (*P*=0.852, *Z*=0.187); Ripple occurrence showed a significant change in CA2 (Ctl>DR: *P*=0.032, *Z*=2.147) and CA3 (Ctl>DR: *P*=0.034, *Z*=2.1262) but not in CA1 (*P*=0.529, *Z*=0.6302). All tests performed using two-sided Wilcoxon rank-sum. **f.** Example of averaged ripple-trigged spectrogram recorded in CA1 during Pre- and Post-CNO sessions. CA1 ripple power is synchronized with SWRs peak in both groups, however a power decrease is seen in DREADD mice in the Post-CNO session. Black lines, SWRs peak; white lines, averaged ripple amplitude. All boxplots represent the median (center mark) and the 25th-75th percentiles, with whiskers extending to the most extreme data points without considering outliers, which were also included in statistical analyses; \**P*<0.05; \*\**P*<0.01; \*\*\**P*<0.001; n.s.= not significant

### Acute silencing of CA2 pyramidal cells impacts hippocampal spiking and oscillations

We first examined how the inhibition of CA2 pyramidal cells (PCs) alters basic hippocampal physiology during rest. Comparing the average firing rates of PCs during Pre- and Post-CNO rest sessions, we observed the expected decrease in CA2 in DREADD mice (Ctl: N=6 mice, 9 sessions; DREADD: N=7 mice, 19 sessions; \**P*<0.05, two-way ANOVA, **Fig. 1c, d**), but no significant changes in CA1 and CA3, indicating a significant and specific silencing of CA2 (**Fig. 1c, d; Extended Data Fig. 1c, d**).

Previous work has suggested CA2 has the ability to trigger SWRs(Boehringer et al., 2017; Oliva et al., 2016a; Oliva et al., 2020), thus we next asked if acute inhibition of CA2 alters the properties or timing of SWRs events. We first examined SWRs in each CA region independently (intra-region), comparing how their properties changed between the first rest session and the last rest session of the day. It has been suggested that spatial experience can lead to increases in SWR power, length and occurrence in subsequent rest sessions (Eschenko et al., 2008; Grosmark and Buzsaki, 2016; Grosmark, 2020; Ramadan et al., 2009); consistent with this, the normalized change in CA1 ripple-band power between the Pre- and Post-CNO sessions increased in control mice, but not in the DREADD mice. However, SWR frequency, duration, and occurrence were similar in both groups across the sessions (**Fig. 1e**). As expected, CA2 SWRs in DREADD mice showed a significant decrease in power, frequency, and occurrence, although the remaining events were significantly longer in duration compared with controls. In CA3 the normalized change in the power, frequency, and duration of SWRs between sessions was similar across the groups, however we found a significantly larger increase in the occurrence of SWRs in control compared to DREADD mice. Accordingly, peri-ripple spectrograms using ripples detected in each region individually as a temporal reference revealed a clear power decrease in CA2 but not for CA3 (**Extended Data Fig. 2a**).

As the majority of SWRs recruit high-frequency activity in all CA regions nearly simultaneously(Buzsaki, 2015; Oliva et al., 2016a; Stark et al., 2014; Sullivan et al., 2011), we next computed wavelet spectrograms of LFP recorded in each region, centered at the peak ripple-band power (100-250 Hz) of events recorded in CA1. Consistent with prior evidence(Fernandez-Lamo et al., 2019; Oliva et al., 2016a), in control mice ripple power in CA1 was strongly aligned to the SWR peak (**Fig. 1f**). However, ripple power decreased following CNO in the DREADD mice in CA1 (**Fig. 1f**), as well as in CA2 and CA3 (**Extended Data Fig. 2b**). Further, cross-correlation analysis of the timing of peak SWR power in CA1 and peaks in CA2 and CA3 also revealed a poorer correlation in DREADD mice compared to controls (**Extended Data Fig. 2c, d**). In summary, DREADD-mediated inhibition significantly reduced CA2 firing, however SWRs were still present, albeit at reduced power in CA1, and with less temporal precision across the CA regions.

### Temporal coordination of spiking activity during SWRs is disrupted in DREADD mice

We next examined spike-LFP dynamics to determine the temporal coordination of spiking during SWRs. Taking the advantage of the DREADD system, which allowed us to compare the same neuronal ensembles before and after CA2 inhibition, we extracted spikes of all PCs in a peri-SWRs window (+/− 100 ms) and calculated a normalized firing curve for each session (Pre-CNO, Post-CNO) in the two groups. In agreement with previous findings(Csicsvari et al., 2000; Kay et al., 2016; Oliva et al., 2016a; Oliva et al., 2020; Sullivan et al., 2011), in control mice CA1 and CA3 neurons discharged close to the CA1 SWRs peak, whereas many CA2 PCs fired earlier and were suppressed during the CA1 and CA3 population discharge (**Fig. 2a, 2e**). In the DREADD mice, CA1 and CA3 PCs firing remained aligned to the CA1 SWR-peak, but a significant decrease in peri-SWR firing rate was found in the Post-CNO compared to the Pre-CNO session. CA2 PCs in DREADD mice displayed a similar ramping trend to controls in the Pre session, but this was lost following CNO-mediated CA2 inhibition (**Fig. 2b, 2e**). This change in firing rate was accompanied by a significant decrease in the participation of CA1, CA2, and CA3 PCs in SWRs during the Post session in DREADD mice (**Fig. 2f**), as well as significantly lower SWR-related bursting in CA1 and CA3 (**Fig. 2g)** and a loss of the experience-dependent increase in recruitment after new learning in CA1 and CA3 (**Extended Data Fig. 3a-c**).

**Figure 2.**
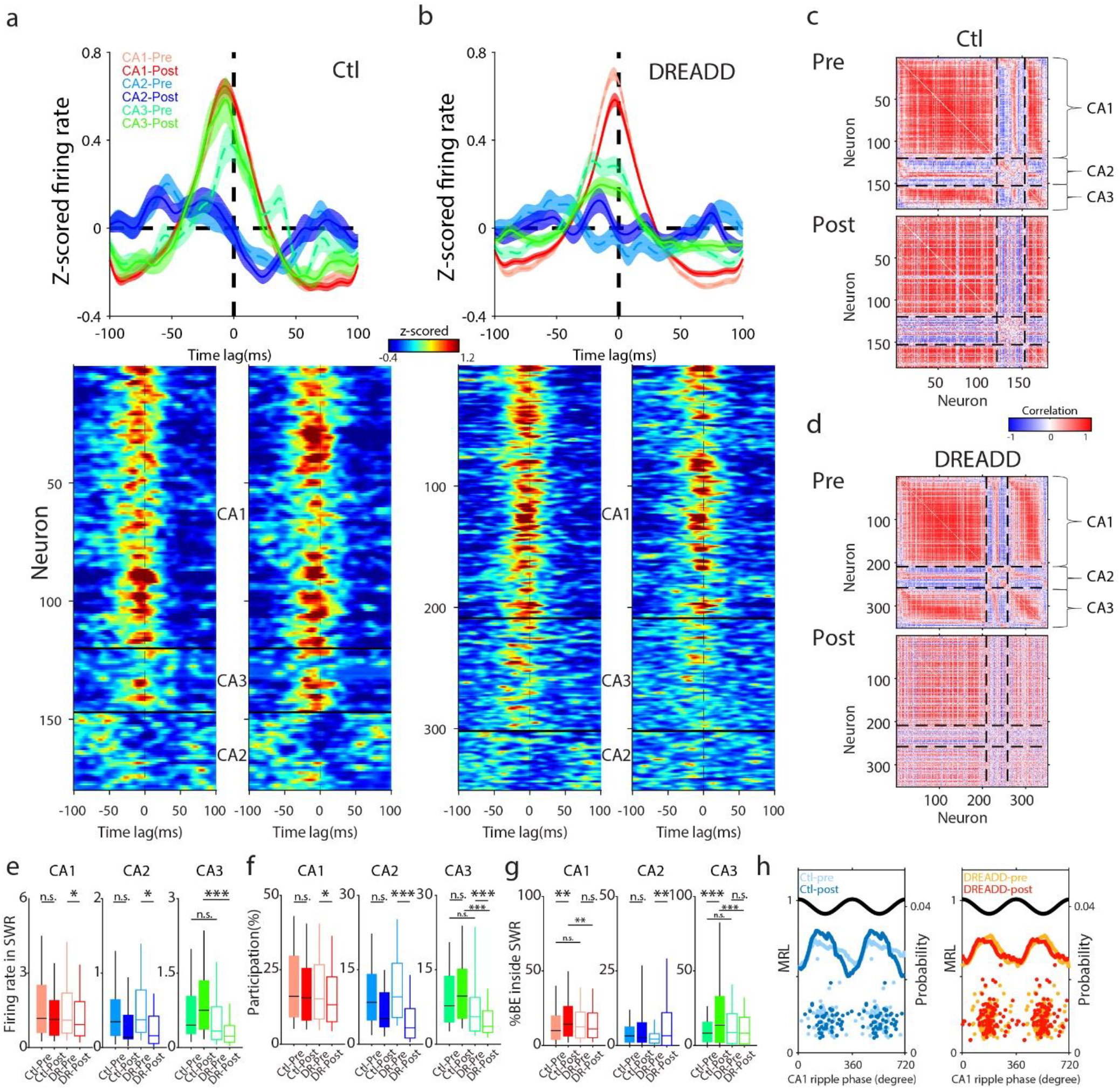
The temporal organization of spiking during SWRs is disrupted following CA2 inhibition. **a.** (upper) Population averaged peri-SWRs z-scored firing rate curves (mean ± sem) and (lower) corresponding z-scored firing rate plots for all pyramidal cells recorded in each CA site for Control mice (CA1: *N*=120 cells; CA3: *N*=27 cells; CA2: *N*=33 cells). Cell IDs were matched between Pre-(firing rate plot, left panel) and Post-CNO (firing rate plot, right panel) session. **b.** Same as in **a.**, but for DREADD mice (CA1: *N*=209 cells; CA3: *N*=93 cells; CA2: *N*=49 cells). Note a strong disruption of the firing pattern of CA2 PCs in the Post-CNO session. **c, d.** Correlation matrix of all individual neuronal peri-SWRs firing curves from a. and b. **c.** Pair-wise correlations of all pooled data for Control mice, top from the Pre-CNO session, bottom from the Post-CNO session.**d**. Same for neurons recorded in the DREADD mice. Cell IDs were matched between Pre- and Post-CNO session for each group. While spatial experiences led to an increase in correlations in the control mice, DREADD mice displayed a loss of temporal coordination accompanied with decreased correlation in Post-CNO session (see **Extended Data 4a-d** for details). **e.** Mean FR of all pyramidal cells during CA1 SWRs. In CA1, a significant main effect of session was found (*F*(1,1051)=7.679, *P*=0.006), with post-hoc comparisons revealing a significant decrease in DREADD mice (pre>post, *P*=0.025) but not in Controls (pre vs. post, *P*=0.072). In CA2, we also observed a significant main effect of session (*F*(1,178)=5.223, *P*=0.023) with post-hoc revealing a significant decrease in DREADD (pre>post, *P*=0.027) but not in Controls (pre vs. post, *P*=0.247). In CA3, a main effect of group was observed (*F*(1,426)=11.274, *P*=0.001), post-hoc comparisons revealed a significant difference in Post-CNO session (Ctl>DR: *P*=0.001) but not in Pre-CNO session (Ctl vs. DR, *P*=0.213). (Two-way ANOVA followed by Bonferroni’s post-hoc, see **Extended Data Table 1** for detailed statistics). **f.** Participation of all pyramidal cells in CA1 SWRs. In CA1, a two-way ANOVA revealed a significant main effect of the session (*F*(1,1051)=4.709, *P*=0.030) and post-hoc tests indicated a significant decrease in DREADD mice (pre>post, *P*=0.037) but not in Controls (pre vs. post, *P*=0.244). In CA2, the main effect was found only in session (*F*(1,209)=16.099, *P*<0.001), with post-hoc showing a significant decrease in DREADD (pre>post, *P*<0.001) but not in Control mice (pre vs. post, *P*=0.103). In CA3, a significant interaction between group and session was found (*F*(1,438)=7.456, *P*=0.007) and post-hoc testing found a significant decrease in DREADD (pre>post, *P*=0.001) but not in Controls (pre vs. post, *P*=0.237), a significant decrease in Post-CNO session (Ctl>DR, *P*<0.001) but not in Pre-CNO session (Ctl vs. DR, *P*=0.114). (Two-way ANOVA followed by Bonferroni’s post-hoc, see **Extended Data Table 1** for detailed statistics). **g.** Percent of burst events (BE) during CA1 SWRs. In CA1, a significant interaction of session and group (*F*(1,1051)=6.476, *P*=0.011) was found, post-hoc revealed a significant increase in Controls (pre<post, *P*=0.005) but not in DREADD (pre vs. post, *P*=0.628) and a significant difference in the Post-CNO session (Ctl>DR, *P*=0.010) but not in Pre-CNO session (Ctl vs. DR, *P*=0.330). In CA2, a main effect was found only in session (*F*(1,209)=11.986, *P*=0.001), post-hoc showed a significant increase in DREADD (pre<post, *P*=0.001) but not in Controls (pre vs. post, *P*=0.066). In CA3, a main effect of the group x session interaction (*F*(1,438)=17.936, *P*<0.001) was observed. Post-hoc comparisons showed a significant increase in Controls (pre<post, *P*<0.001)b ut not in DREADD (pre vs. post, *P*=0.222), a significant difference was found also in Post-CNO session (Ctl>DR, *P*<0.001) but not in Pre-CNO (Ctl vs. DR, *P*=0.123). (Two-way ANOVA followed by Bonferroni’s post-hoc, see **Extended Data Table 1** for detailed statistics). **h.** Ripple phase preference was similar between groups in both strength (*P*=0.608, *F*=0.611, ANOVA) and preferred phase (*P*=0.429, *F*=0.925, circular ANOVA). Averaged phase probabilities (solid line) are shown for all significantly modulated cells (*P*<0.05, Rayleigh test), scatter plots indicate the preferred phase and mean resultant length (MRL) for all significantly modulated cells (see **Extended Data Fig. 4e-f** for details). All boxplots represent the median (center mark) and the 25th-75th percentiles, with whiskers extending to the most extreme data points without considering outliers, which were also included in statistical analyses; \**P*<0.05; \*\**P*<0.01; \*\*\**P*<0.001; n.s.= not significant

To examine how these changes in firing altered interactions between neurons, we next calculated pairwise correlations of the normalized peri-SWRs firing rate curve of each neuron for each session in the two groups. Following spatial experience a more structured and clustered pattern emerged in the controls during the Post-CNO compared to the Pre-CNO sessions, consistent with previous data finding experience-dependent changes in coordinated activity(Grosmark and Buzsaki, 2016; Grosmark, 2020; Wilson and McNaughton, 1994) (**Fig. 2c; Extended Data Fig. 4a-d**). However, in the DREADD group the structured pattern was lost, both within and across CA regions, following CA2 inhibition, reflected as a significantly lower average correlation value across all pairs (Pearson’s r) (**Fig. 2d; Extended Data Fig. 4a-d**). Despite these changes, the ripple phase preference was comparable between groups in both strength and preferred angle (**Fig. 2h; Extended Data Fig. 4e, f**). These data suggest that the experience-dependent facilitation of CA1 and CA3 spike recruitment and the resulting ensemble interactions during SWRs may be lost following CA2 silencing.

### Temporal precision of cell assembly reactivation during SWRs is reduced following inhibition of CA2

Given the loss of pairwise coordination observed in single-cell activity during SWRs, we hypothesize that the structured reactivation of neuronal ensembles may also be impacted. To evaluate this possibility a combination of principal component (PCA) and independent component analyses (ICA) was employed to detect the co-activation of cell assemblies in CA1 during exploration and then the re-expression of these assemblies was tracked over time. We first used the place cell activity on LT2 to identify assembly patterns (Ctl: N=6 mice, 7 sessions; DREADD: N=5 mice, 7 sessions). Consistent with previous work(Middleton et al., 2018; van de Ven et al., 2016), the cell assemblies detected were spatially tuned (**Fig 3a; Extended Data Fig. 5a**). The expression strength of the assemblies on LT2 was similar between the groups (**Extended Data Fig. 5f**), consistent with our analysis of place cell rate coding (**Extended Data Fig. 5b-e**). We further confirmed that neurons belonging to the same assembly had stronger co-firing coefficients and place-field similarity than other neuronal pairs (**Extended Data Fig. 5g**) in both groups, indicating a comparable capability of population spatial coding following the silencing of CA2. We next tracked the activity of the cell assemblies identified on LT2 during the following sleep session (Post-CNO). We observed that across the entire session the reactivation rate was comparable between groups (Ctl: 0.155±0.012, DREADD: 0.115±0.012, *P*>0.05, **Fig. 3d**), however the reactivation strength was remarkably higher in DREADD compared to control mice (Ctl: 18.866±1.008, DREADD: 34.760±5.805, \*\*\**P*<0.001, **Fig. 3d**).

**Figure 3.**
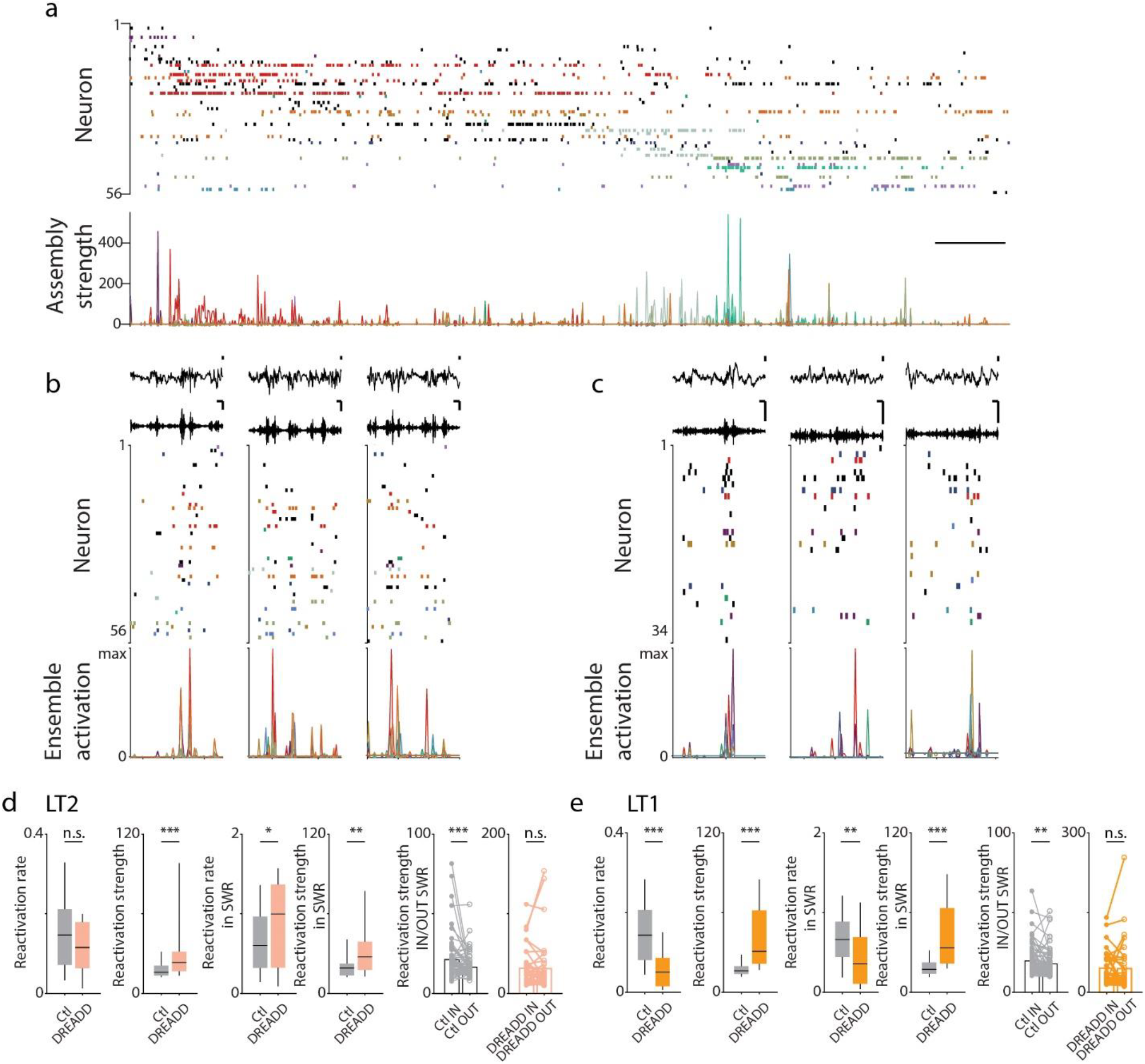
The temporal precision of SWR-associated assembly reactivation is decreased in CA2 DREADD mice. **a.** Example of CA1 neuronal activity during a single lap on LT2 from a control mouse. The raster plot on top indicates the firing of 56 simultaneously recorded pyramidal cells. The colors represent individual cell assemblies (13 cell assembly overall). Spikes in black are shown for completeness but do not belong to any significantly detected cell assemblies. The expression strength of each cell assembly is shown on the bottom. The horizontal scale bar denotes 2 s (see **Extended Data Fig. 5a** for DREADD mouse example). **b.** Examples of reactivation patterns identified from the LT2 during the Post-CNO session for the same Control mouse in **a.** The top panel shows raw and filtered (100-180Hz) CA1 SWR-LFPs. Middle panel shows spiking activity within each color-coded cell assemblies aligned by their positions on the track. The bottom panel shows reactivation strength in the same temporal bins. Scale bars denotes 100 ms (horizontal) and 200 μV (vertical) respectively. **c.** Same as in **b**, but for a DREADD mouse. **d.** Reactivation properties of LT2 assemblies during the Post-CNO session. (From left to right) Reactivation rate was comparable between groups (*P*=0.142, *Z*=1.469, Wilcoxon rank-sum), while DREADD mice displayed a significantly higher reactivation strength than Controls (*P*=5.801×10^−4^, *Z*=-3.441, Wilcoxon rank-sum) throughout the session. The reactivation rate (*P*=0.048, *Z*=-1.979, Wilcoxon rank-sum) and strength (*P*=0.001, *Z*=-3.265, Wilcoxon rank-sum) were both significantly higher for DREADD mice compared to controls during SWRs. **e.** Reactivation strength inside and outside SWRs showed a significant difference in Control mice (IN>OUT, *P*=2.785×10^−4^, *Z*=3.635, Wilcoxon sign-rank), while it was unchanged in DREADD mice (IN vs. OUT, *P*=0.219, *Z*=1.228, Wilcoxon sign-rank). Reactivation properties in Post-CNO session of LT1 assemblies. (From left to right) Reactivation rate was significantly reduced in DREADD mice compared to Controls (*P*=1.350×10^−7^, *Z*=5.2719, Wilcoxon rank-sum), while a significant increase was observed in reactivation strength (*P*=1.282×10^−9^, *Z*=-6.098, Wilcoxon rank-sum) across the session. During SWRs, the reactivation rate (*P*=0.004, *Z*=2.867, Wilcoxon rank-sum) was significantly decreased in DREADD mice compared to Controls, whereas strength (*P*=2.934×10^−9^, *Z*=-5.935, Wilcoxon rank-sum) was significantly higher. Reactivation strength inside and outside SWRs showed a significant difference in Controls (IN>OUT, *P*=0.001, *Z*=3.215, Wilcoxon sign-rank), while it was similar in DREADD mice (IN vs. OUT, *P*=0.116, *Z*=1.571, Wilcoxon sign-rank). All boxplots represent the median (center mark) and the 25th-75th percentiles, with whiskers extending to the most extreme data points without considering outliers, which were also included in statistical analyses; \**P*<0.05; \*\**P*<0.01; \*\*\**P*<0.001; n.s.= not significant

Robust reactivation of cell assemblies occurs during SWRs, a phenomena thought to support memory consolidation(Buzsaki, 2015; van de Ven et al., 2016); thus we compared assembly activity in the groups both inside and outside of SWR-periods (**Fig. 3b, 3c**). We found that DREADD mice showed a significantly higher reactivation rate (Ctl: 0.676±0.051, DREADD: 0.883±0.095, \**P*<0.05, **Fig. 3d**) and strength (Ctl: 21.293±1.545, DREADD: 31.479±3.668, \*\**P*<0.01, **Fig. 3d**) during SWRs compared to controls. Moreover, while reactivation strength in controls was significantly higher within compared to outside SWRs, in the DREADD mice it remained high throughout both periods (DREADD: Inside: 31.479±3.668, Outside: 31.238±5.229, *P*>0.05; Ctl: Inside: 21.293±1.545, Outside: 16.475±0.803, \*\*\**P*<0.001, **Fig. 3d**), indicating a loss of temporal coordination with SWR events.

Given that the LT2 assemblies were formed following CA2 inhibition, we next asked if the same phenotypes could be seen in assemblies active on LT1, prior to CNO injection. LT1 assemblies were detected and their reactivation was examined in the Post-CNO session (Ctl: N=6 mice, 7 sessions; DREADD: N=5 mice, 7 sessions). We found that LT1 assemblies reactivated less frequently in DREADD compared to control mice across the whole session (Ctl: 0.149±0.010, DREADD: 0.063±0.010, \*\*\**P*<0.001, **Fig. 3e**), as well as during SWR-period (Ctl: 0.690±0.043, DREADD: 0.500±0.078, \*\**P*<0.01, **Fig. 3e**). However, similar to LT2 assemblies, reactivation strength was significantly higher in the DREADD group, across the whole session (Ctl: 18.565±0.857, DREADD: 45.751±6.375, \*\*\**P*<0.001, **Fig. 3e**), during SWRs (Ctl: 19.668±1.005, DREADD: 45.360±5.242, \*\*\**P*<0.001, **Fig. 3e**), and outside SWR-periods (DREADD: Inside: 45.360±5.242, Outside: 44.583±7.796, *P*>0.05; Ctl: Inside: 19.668±1.005, Outside: 17.841±0.879, \*\**P*<0.01, **Fig. 3e**). Further, we examined the reactivation pattern of LT1 assemblies immediately following drug injection in the subsequent CNO session, during which CA2 was inhibited over the course of roughly 20 minutes(Boehringer et al., 2017) (**Extended Data Fig. 6a**). While reactivation was comparable between the DREADD and control groups at the start of the session, the increase in reactivation strength, both inside and outside SWRs, emerged 20 minutes after CNO injection (**Extended Data Fig. 6b, c**). Together, these results indicate that the SWR-associated temporal precision of offline assembly reactivation is impaired in the absence of CA2 transmission and CA2 inhibition results in increase in the strength of reactivation of recently encoded cell assemblies.

### Diminished specificity of assembly reactivation is accompanied by poorer replay content following silencing of CA2

The increase in reactivation strength and the decrease in coordination between reactivation and SWR events of both LT1 and LT2 assemblies in the DREADD mice motivated us to next assess the temporal relationship of the reactivation of the distinct LT1 and LT2 cell assemblies during the Post-CNO session. We first examined the cross-correlation between assembly event-time and SWR-onset time and observed a comparable peak correlation between groups both for LT1 (Ctl: 0.079±0.003, DREADD: 0.089±0.007, *P*>0.05, **Fig. 4a**) and LT2 assemblies (Ctl: 0.082±0.003, DREADD: 0.088±0.004, *P*>0.05, **Fig. 4a**). However, fine scale temporal analysis revealed a significant leftward shift of the peak activation relative to SWR-onset in the DREADD mice both for LT1 (Ctl: −0.012±0.001s, DREADD: −0.022±0.004s, \**P*<0.05, **Fig. 4a**) and LT2 assemblies (Ctl: −0.010±0.001s, DREADD: −0.022±0.002s, \*\*\**P*<0.001, **Fig. 4a**) compared with controls. A similar result was obtained when we compared the reactivation strength within-SWR-period with the pre-SWR-period, again for both LT1 and LT2 assemblies (**Extended Data Fig. 7a, b**).

**Figure 4.**
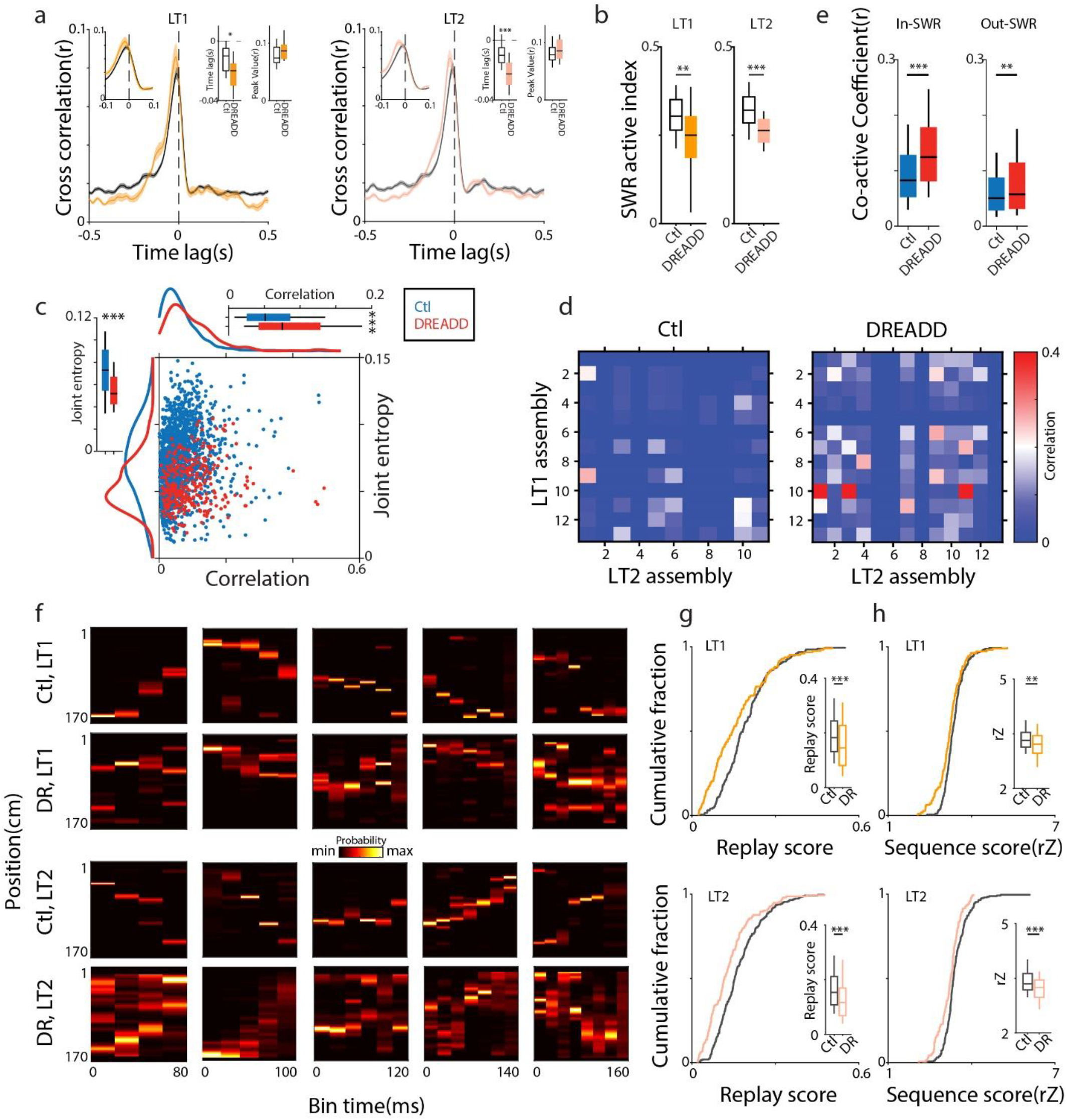
CA2 DREADD mice exhibit reduced assembly specificity and poorer replay quality. **a.** Averaged cross-correlation between assembly event and SWR-onset time. Insets provide a zoomed (−0.1s to 0.1s; left) view around the zero-lag time point and quantification (right). Assembly events in DREADD mice reactivated significantly earlier in relation to ripple-onset than Controls. Left (LT1): time lag (*P*=0.015, *Z*=2.446), peak value (*P*=0.171, *Z*=-1.369); Right (LT2): time lag (*P*=2.763×10^−5^, *Z*=4.192), peak value (*P*=0.190, *Z*=-1.312). Two-sided Wilcoxon rank-sum. Line corresponds to mean, shaded region corresponds to s.e.m.. **b.** SWR active-index demonstrated a reduction of SWR-modulation for DREADD mice both for LT1 (Ctl>DR: *P*=0.002, *Z*=3.142) and LT2 (Ctl>DR: *P*=7.018×10^−5^, *Z*=3.976) assemblies compared with Controls. Two-sided Wilcoxon rank-sum. **c.** Scatter plot with density histograms showing the coordination between LT1 and LT2 assembly events throughout the Post-CNO session for two groups. DREADD mice showed a significant increase in correlation (Ctl<DR: *P*=1.585×10^−13^, *Z*=-7.380) but a decrease in joint entropy (Ctl>DR: *P*=1.412×10^−29^, *Z*=11.2935), indicating a reduction in the specificity and independency of two representations. Two-sided Wilcoxon rank-sum. **d.** Example of the correlation matrix between LT1 and LT2 assemblies during Post-CNO session for two groups. Correlations were significantly higher in CA2 DREADD mice. **e.** Correlation of LT1 and LT2 assembly in/out SWRs during Post-CNO session. Group data displayed a significantly higher correlation inside-SWR (left panel) (Ctl<DR: *P*=6.440×10^−22^, *Z*=-9.622) and also outside-SWR (right panel) (Ctl<DR: *P*=0.002, *Z*=-3.123) in DREADD compared to Controls. Two-sided Wilcoxon rank-sum. **f.** Examples of replay sequences derived from Bayesian decoding during SWRs in Post-CNO session. Note poorer replay content was observed (for both LT1 and LT2) in DREADD mice compared to Controls. **g.** Quantification of replay sequences by replay score. Cumulative distributions of replay score for LT1 (top) and LT2 (bottom), with medians and data distribution shown in the inset. DREADD mice displayed significantly reduced scores for LT1 (Ctl>DR: *P*=6.799×10^−4^, *Z*=3.398) and LT2 (Ctl>DR: *P*=6.602×10^−7^, *Z*=4.973) replays compared with Controls. Two-sided Wilcoxon rank-sum. **h.** Quantification of replay sequences by sequence score (rZ). Cumulative distributions of rZ for LT1 (top) and LT2 (bottom), with medians and data distribution shown in the inset. DREADD mice displayed significantly reduced scores for both LT1 (Ctl>DR: *P*=0.003, *Z*=2.953) and LT2 (Ctl>DR: *P*=2.867×10^−5^, *Z*=4.184) sequences compared with Controls. Two-sided Wilcoxon rank-sum. All boxplots represent the median (center mark) and the 25th-75th percentiles, with whiskers extending to the most extreme data points without considering outliers, which were also included in statistical analyses; \**P*<0.05; \*\**P*<0.01; \*\*\**P*<0.001; n.s.= not significant

We next calculated a SWR active-index (see Methods) to quantify how strongly reactivation events were modulated by SWRs and observed a significant decrease in DREADD mice for both LT1 (Ctl: 0.307±0.008, DREADD: 0.247±0.023, \*\**P*<0.01) and LT2 assemblies (Ctl: 0.321±0.010, DREADD: 0.261±0.009, \*\*\**P*<0.001, **Fig. 4b; Extended Data Fig. 7c, d**). This clear reduction of SWR-modulation in DREADD mice raises the question of whether the two distinct assembly representations (LT1, LT2) are maintained during the Post-CNO session. To answer that, we classified individual SWRs into 3 categories (LT1-assembly only, LT2-assembly only, LT1 and LT2 assembly both) and calculated the fraction of SWRs for each category (**Extended Data Fig. 8a**). DREADD mice showed an asymmetrical difference compared to control mice, with a decrease in LT1 SWRs and an increase in LT2 SWRs, as well as a significantly higher fraction of SWRs containing both LT1 and LT2 assemblies (LT1 ∩ LT2: Ctl: 0.009±0.0002, DREADD: 0.014±0.0006, \*\*\**P*<0.001, **Extended Data Fig. 8b**). A co-active strength (see Methods) of LT1 and LT2 assemblies also revealed a significantly higher value in DREADD mice (Ctl: 0.039±0.0004, DREADD: 0.055±0.001, \*\*\**P*<0.001, **Extended Data Fig. 8c**).

To quantify event synchrony we employed information theory and found a significantly higher correlation (Ctl: 0.064±0.001, DREADD: 0.093±0.004, \*\*\**P*<0.001, **Fig. 4c, 4d**) and a significantly lower joint entropy (Ctl: 0.072±0.0006, DREADD: 0.055±0.0009, \*\*\**P*<0.001, **Fig. 4c**) between LT1 and LT2 assemblies in DREADD mice compared with controls, results that were further confirmed by higher normalized mutual information (Ctl: 0.024±0.0007, DREADD: 0.042±0.003, \*\*\**P*<0.001) and lower conditional entropy (Ctl: 0.033±0.0004, DREADD: 0.020±0.0007, \*\*\**P*<0.001, **Extended Data Fig. 8d**). The higher correlation of the LT1 and LT2 assemblies in the DREADD mice was observed both within and outside SWR-periods (In-SWR: Ctl: 0.095±0.002, DREADD: 0.138±0.004, \*\*\**P*<0.001; Out-SWR: Ctl: 0.065±0.002, DREADD: 0.083±0.005, \*\**P*<0.01, **Fig. 4e**). This is also consistent with a higher pair-wise correlation for distinct LT1-LT1 and distinct LT2-LT2 assembly pairs in the DREADD mice (LT1: Ctl_whole-session:_ 0.073±0.002, DREADD_whole-session_: 0.104±0.008, \*\**P*<0.01; Ctl_within-SWRs_: 0.095±0.002, DREADD_within-SWRs_: 0.138±0.009, \*\*\**P*<0.001; Ctl_out-SWRs_: 0.064±0.002, DREADD_out-SWRs_: 0.082±0.009, *P*>0.05; LT2: Ctl_whole-session_: 0.087±0.003, DREADD_whole-session_: 0.151±0.008, \*\*\**P*<0.001, Ctl_within-SWRs_: 0.106±0.004, DREADD_within-SWRs_: 0.189±0.008, \*\*\**P*<0.001; Ctl_out-SWRs_: 0.080±0.003, DREADD_out-SWRs_: 0.124±0.009, \*\*\**P*<0.001, **Extended Data Fig. 8e, f**). These findings indicate a higher similarity and lower independence of the two representations reactivated in the time domain and results in a loss of informational precision following the silencing of CA2.

Finally, to determine the effect of this increase in joint activation on sequential activation, place cell firing rate curves during LT1 and LT2 track exploration were constructed to decode sequential trajectories (‘replays’) during SWRs in the Post-CNO session. Consistent with prior work(Silva et al., 2015), independent and structured replay sequences were observed both for LT1 and LT2 in control mice (N=6 mice, 7 sessions; **Fig. 4f**). However, DREADD mice (N=5 mice, 7 sessions) displayed poorer and noisier replay content (**Fig. 4f**), which following decoding appeared as repeated vertical “stripes” of activity across time. To statistically quantify the quality of trajectory events for the different tracks, a replay score(Davidson et al., 2009), which represents the estimated likelihood of an experienced trajectory (see Methods), was calculated. We found significantly lower replay scores for both LT1 and LT2 in DREADD compared with control mice (LT1: Ctl: 0.195±0.006, DREADD: 0.145±0.009, \*\*\**P*<0.001; LT2: Ctl: 0.169±0.005, DREADD: 0.132±0.007, \*\*\**P*<0.001, **Fig. 4g**). We then calculated a normalized Z-score value (rZ sequence score(Grosmark and Buzsaki, 2016)) to quantify significance of individual replay events. This analysis also revealed significantly lower scores for the two tracks in DREADD compared with control mice (LT1: Ctl: 3.374±0.027, DREADD: 3.222±0.043, \*\**P*<0.01; LT2: Ctl: 3.425±0.027, DREADD: 3.209±0.032, \*\*\**P*<0.001, **Fig. 4h**). Together, these results demonstrate a disruption of the replay structure in the DREADD group.

The fact that replay sequences still occurred, but were less sharply tuned in DREADD mice (sharpness: LT1: Ctl: 0.272±0.008, DREADD: 0.241±0.015, \*\*\**P*<0.001; LT2: Ctl: 0.222±0.008, DREADD: 0.145±0.008, \*\*\**P*<0.001, **Extended Data Fig. 9e, f**), led us to consider how this may be linked with the altered assembly reactivation described above. As can be seen in a combined frame (**Extended Data Fig. 9a-d**), DREADD mice showed a much shorter replay length and much stronger population synchrony within each time bin compared to controls. To quantify this, we first calculated replay distance, which showed a significant decrease for both LT1 and LT2 in DREADD mice compared to controls (LT1: Ctl: 90.939±4.365, DREADD: 53.266±4.023, \*\*\**P*<0.001; LT2: Ctl: 83.091±3.833, DREADD: 68.747±4.382, \**P*<0.05, **Extended Data Fig. 9e, f**). Next, a normalized max participation index (see Methods) was calculated, representing the degree of population discharge for each candidate replay. This analysis revealed a significantly higher value for both LT1 and LT2 in DREADD mice (LT1: Ctl: 0.087±0.003, DREADD: 0.142±0.007, \*\*\**P*<0.001; LT2: Ctl: 0.104±0.003, DREADD: 0.128±0.005, \*\*\**P*<0.001, **Extended Data Fig. 9e, f**), suggesting higher population synchrony during each time bin in DREADD mice, resulting in shortened replay span and the “stripe” like replay structure. This observation is consistent with the elevated assembly reactivation strength we observed in the DREADD mice. Though LT1 assemblies were formed with CA2 intact (prior to CNO), the DREADD group displayed a generalized disruption of replay content for both tracks, suggesting ongoing CA2 activity is required for the precise SWR-associated reactivation of previously formed memories.

## Discussion

Here we find that acute DREADD-mediating silencing of CA2 leads to changes in both SWRs and the underlying structured patterns of CA1 spiking linked to the consolidation of memory. While we did not observe a change in the number of SWR events following CA2 inhibition(Alexander et al., 2018), we did see a reduction in ripple band power in CA1 (**Fig. 1e-f**). Given previous evidence that CA2 can generate ripples independent of CA3(Oliva et al., 2016a), the lack of change in SWR number suggests that even acute silencing of CA2 may result in a compensatory increase in CA3 driven SWRs, perhaps due to a decrease in CA2-mediated feedforward inhibition(Boehringer et al., 2017). More strikingly, the temporal precision of SWR activity between CA1 and CA3 was reduced in the absence of CA2 transmission (**Extended Data Fig. 2b-d**), suggesting a role in coordinating synchrony between the regions, despite their direct connectivity.

On the level of neuronal activity, acute CA2 silencing had little impact on general pyramidal cell activity in CA1 and CA3, both during novel track exploration and rest, yet had a profound effect on the experience-dependent increase in SWR-related spiking, both within and across these regions (**Fig. 2c, 2d; Extended Data Fig. 3a-c, 4a-d**). This was even more pronounced on the level of neuronal assemblies. The loss of CA2 output increased the strength of reactivation of CA1 assemblies, consistent with a role in mediating inhibition in the neighboring CA regions(Boehringer et al., 2017). Despite this increase in strength, there was a profound loss of temporal precision, manifest as increased assembly reactivation outside of SWRs and less fine temporal precision around the SWR event itself (**Fig. 3d-e; Fig. 4a; Extended Data Fig. 7a-b**). This was true for assemblies formed both prior to (LT1) and following (LT2) CA2 inhibition, suggesting this is not a state dependent effect and reflects a role for CA2 in shaping both the timing and content of reactivation events.

The increase in reactivation strength of the experience-dependent assemblies also resulted in the significant increase in the co-activation of LT1 and LT2 assemblies. Prior work has shown that reactivation of multiple distinct experiences can occur during rest(O’Neill et al., 2010; Silva et al., 2015), with orthogonal representations temporally segregated to distinct SWR events. However, following CA2 inhibition the inappropriate co-activation of assemblies from distinct experiences rose significantly (**Fig. 4c-e; Extended Data Fig. 8a-d**), which may lead to interference between these memories. These finding suggest a key contribution of ripple-associated CA2 activity may be in the selection or prioritization of specific assemblies to participate in a given SWR event(Stober et al., 2020). Whether this is due to CA2 input directly into CA1 or its recruitment of local inhibition in CA3(Boehringer et al., 2017) cannot be determined from the current data. However, given that chronic silencing of CA2 transmission results in an increase in excitability in the CA3 recurrent network(Boehringer et al., 2017), acute silencing may have a similar, although subtler, impact, leading to increase in both reactivation strength of single assemblies and co-activation of assemblies representing distinct experiences.

Finally, analysis of replay events revealed an increased participation and synchrony of CA1 neurons, resulting in a decrease in the quality and length of these events crucial for memory consolidation. This was true for the replay of place cell sequences from both LT1, experienced with CA2 activity intact, and LT2, experienced after CA2 inhibition. These data suggest that CA2 activity is required for both the selection of the informational template to be re-expressed and the temporal precision of the reactivation itself. The precise temporal activity of CA2 neurons in relation to the SWR onset in these processes cannot be determined with DREADD-mediating silencing. It has been suggested that CA2 activity during periods of non-movement outside of ripples(Kay et al., 2016) could play a role in influencing the reactivation and abstraction of previous experiences(Kay and Frank, 2019). Future experiments with increased temporal and cell-type specificity will be necessary to test these hypotheses.

While much of the CA2 literature has focused on its preferential role in social behavior(Hitti and Siegelbaum, 2014; Oliva et al., 2020; Piskorowski et al., 2016; Smith et al., 2016; Young et al., 2006), our findings point to a more general role in coordinating both the temporal and informational precision of activity during SWRs, even following purely spatial experiences. As has been suggested, CA2 activity may contribute to the excitatory drive needed to generate SWRs(Oliva et al., 2016b), either alone or in concert with CA3. This is consistent with the decrease in average ripple band power we observed in CA1 in the DREADD mice (**Fig. 1e-f**). However, our data suggest CA2 may play an additional role in coordinating and filtering spiking activity within and between CA3 and CA1. This may involve regulating a precise inhibitory/excitatory balance onto specific neuronal assemblies in these regions to shape which neurons participate in a given SWR (information), as well as the temporal dynamics of this activation. Thus CA2 may both assist in the general excitatory drive required for SWR generation, while providing input to bias the informational content of the replay events themselves.

## Methods

### Subjects

All procedures were approved by the RIKEN Animal Care and Use Committee. All experiments were conducted with a cohort of male mice hemizygous for the Cacng5-Cre transgene (age 3-5 months) injected with Cre-dependent AAV virus into CA2 that were previously described(Boehringer et al., 2017). The mice were housed in transparent Plexiglas cages placed in ventilated racks and maintained in a temperature- and humidity-controlled room with a 12-h light/dark cycle (lights on from 08:00 to 20:00). Prior to stereotaxic surgery all animals were housed in groups of 2 to 5 and provided with food and water *ad libitum.* Following microdrive implantation the mice were single housed. Prior to *in vivo* electrophysiology experiments the animals were experimentally naïve and randomly assigned in respect to which virus was injected. All experiments were conducted during the animals’ light cycle.

### Adeno-associated viruses (AAV) vector construction and production

The pAAV.hSyn.DIO.hM4D.Gi.mCherry plasmid was a gift from Bryan Roth (Addgene plasmid # 44362) and the recombinant AAV vectors pAAV.EF1a.DIO.mCherry, was generated within our laboratory as previously reported(Boehringer et al., 2017). For adeno-associated virus production, we used the AAV Helper Free System (Agilent Technologies). The adeno-associated virus vector (pAAV.EF1a.DIO.mCherry or pAAV.hSyn.DIO.hM4D.Gi.mCherry) was co-transfected with pAAV-DJ/8 (Cell Biolabs), which supplies AAV2 replication proteins and AAV-DJ/8 capsid proteins, and pHelper (Agilent Technologies) which supplies the necessary adenovirus gene products required for the AAV production into the 293FT cell line (Invitrogen) utilizing the 293fectin transfection reagent (Invitrogen). After 72 hours, the supernatant was collected and centrifuged at 3,000 rpm for 30 minutes and then filtered through a 0.45μM filtration unit (Millipore). Purification of the AAV was carried out by ultracentrifugation (87,000 g, 4°C, 2 h) with a 20 % sucrose cushion. After ultracentrifugation, the supernatant was removed and the pellet was resuspended in phosphate-buffered saline (PBS), aliquoted and stored at −80°C for long term storage. The AAV stocks were titered using a custom ordered AAV stock purchased from Virovek (Hayward, CA) as the reference standard. AAV titer quantification was performed using qPCR with the StepOne Plus Real Time PCR System (Applied Biosystems), FastStart Universal SYBR Green Master (Roche, Basel), and QPCR primers for an approximately 100 bp fragment of the woodchuck hepatitis virus posttranscriptional response element (WPRE) found in all our adeno-associated virus vectors. The titers for the AAV stocks ranged between 1.37 × 10^12^ to 3.34 × 10^13^ viral genome (vg)/mL.

### Tetrode drives

Custom microdrives were manufactured with the assistance of the Advanced Manufacturing Support Team, RIKEN Center for Advanced Photonics, Japan. The drives contain eight independently adjustable nichrome tetrodes (14μm) (TT), gold-plated to an impedance of 200 to 250 kΩ, arranged in two rows of 4 TT, each of them running along the CA3 to CA1 axis of the dorsal hippocampus.

### Injection of AAV into CA2 and tetrode drive implantations

Mice were anesthetized using Avertin (2, 2, 2-tribromoethanol; Sigma-Aldrich, 476 mg/kg, i.p) and placed into a stereotactic frame (Kopf). AAV was microinjected bilaterally into CA2 (coordinates from bregma: AP −1.5 mm; ML +/−1.8: DV −1.5; 500 nL/hemisphere; injection speed: 100 nL/min; post injection waiting period: 10 min). Control mice (N=6) were injected with AAV.DJ/8.DIO.EF1a.mCherry (9 × 10^8^ vg) and CA2-DREADD (N=7) mice were injected with AAV.DJ/8.hSyn.DIO.hM4D.Gi.mCherry (9 × 10^8^ vg). The recording drive was placed over the injection site in the right dorsal hippocampus with recording tetrodes extending approximately 1 mm.

### Experimental protocol and recording procedures

Over several days the tetrodes were slowly lowered to CA1, CA2, and CA3 *stratum pyramidale* while the animal was maintained in a small sleep box (15 cm diameter × 30 cm high). Approximately two weeks following implantation recording commenced once stable putative cell clusters and hippocampal EEG patterns (sharp-wave ripples) were observed. At the start of every recording day the tetrodes were finely adjusted to maximize cell yield. Both control and DREADD mice underwent multiple days of recording and CNO injection and on every recording day there was a Pre-CNO and a Post-CNO rest session, allowing us to determine the impact of DREADD activation on neuronal activity and hippocampal oscillations as shown in Figs. 1 and 2 (Ctl: N=6 mice, 9 total sessions; DREADD: N=7 mice, 19 total sessions). The data used to analyze assembly activity and replay (Figs. 3 and 4) were collected in 7 sessions from 6 control mice and in 7 sessions from 5 DREADD mice. Those sessions began with a 30 min recording in a small familiar sleep box (Pre-CNO session), which was followed by exploration on a linear track (LT1, 170 L × 10W × 12 H cm; smooth black floor, clear walls; cleaned with 70% ethanol) lasting either 10 min or until 15 laps had been completed. Immediately after, the mice received an intraperitoneal injection of 2mg/kg clozapine-N-oxide (CNO) and remained in the sleep box for 50 min (CNO session), while changes in CA2 spiking were monitored. Consistent with previous finding(Alexander et al., 2018; Boehringer et al., 2017), inhibitory action of the hM4Gi-DREADD receptor was observed to reach full efficacy about 20 min post-injection. Following CNO administration, mice were exposed to a novel linear track (LT2, 170 L × 17 W × 18 H cm; textured brown floor and striped walls; cleaned with 0.5% isoamyl acetate in 70% ethanol). The recoding day ended with a final 40-minute rest session in the same sleep box (Post-CNO session). Recording stability was confirmed by comparing activity during the rest sessions.

During recording, spike data, local field potential (LFP) and behavioral position data were collected using a 32-channel Digital Lynx 4S acquisition system (Neuralynx). The LFP was filtered at 1-9000 Hz. The spike waveforms sampled at 32,556 Hz were filtered between 0.6 and 6 kHz with a peak threshold over 50 μV or 10% of the gain, whichever was larger. Light-emitting diodes (red and green) on the recording headstage were video tracked at 30 Hz to obtain the animal’s position and head direction. At the conclusion of the experiment, mice were anesthetized and electrode positions were marked by electrolytic lesions (with 50 μA current for 8 s through each tetrode individually). Transcardial perfusion with 4% paraformaldehyde (PFA) followed, after which brains were removed and postfixed for a further 24 h in 4% PFA. Coronal slices 50 μm thick were prepared on a vibratome (Leica) and inspected by fluorescent microscopy to confirm electrode placement in relation to CA2 mCherry expression.

### Spike sorting and unit classification

Spikes were manually sorted using SpikeSort3D software (Neuralynx), with putative units clustered in three-dimensional projections of multidimensional parameter space (using both waveform amplitude and energy). To ensure tetrode stability, units were clustered in the Pre-CNO session and cluster boundaries were transpose onto the latter session (LT1/CNO/LT2/Post-CNO) individually; clusters whose spikes fell outside the original bounds were excluded from further analyses. Additionally, clusters were excluded from further analyses if clusters contained > 0.5% spikes having inter-spike-interval of < 2 ms, a total number of spikes < 50 or an isolation distance(Schmitzer-Torbert et al., 2005) < 10. Remaining units were considered pyramidal neurons if mean spike width exceeded 200 μs and had a complex spike index (CSI(McHugh et al., 1996)) ≥5. Clustered units were matched across sessions as previously described(Tanaka et al., 2018), all subsequent analyses were performed in MATLAB (MathWorks, R2017b), using custom written scripts.

### Ripple analyses

Ripple events were detected using methods similar to those previously described(Boehringer et al., 2017). The wide band LFP were firstly band-pass filtered between 100 and 250 Hz using 69 order Kaiser-window FIR zero-phase shift filter. The absolute value of Hilbert transform was then smoothed with 50 ms Gaussian window and candidate ripple events were detected as periods where magnitude exceeded 3SD above the mean for at least 30 ms. The start and end periods of each candidate event were defined as periods when the magnitude returned to the mean. To rule out the false detection, strict criteria were applied. First, the multi-unit activity (MUA) recorded from the same tetrode as the LFP was converted to instantaneous firing rate and smoothed, to allow detection of firing bursts using the same thresholds as described for LFP; any candidate ripple events not coincident with MUA bursts was excluded from subsequent analysis. For cleaner detection of ripple frequency for each ripple, a multitaper method was performed on the product of each filtered ripple waveform and a Hamming window of the same length. To calculate the ripple-triggered spectrograms, the complex wavelet transform (Morlet wavelet, parameter = 7) was first applied to the wide band LFP. Wavelet spectrograms were then calculated for each detected ripple in 400 ms window centered on each ripple. Resulting spectrograms were averaged across individual events to construct final plots.

### Local field potential analyses

Spectral power of LFP was estimated by using pwelch function in the Matlab with 2048 samples window size (1.26 s), 50% overlap and 4096 FFT points (2.52 s). Power was calculated across the frequency range (1-500 Hz). To account for power fluctuations caused by difference in position/impedance of the electrodes and to make power values comparable across mice, a single electrode from each animal situated in CA1 *stratum pyramidale* was used for analysis in both the Pre- and Post-CNO session and values were normalized as the ratio (Post/Pre). For spike-LFP phase analysis, phase preference for ripples (100-180 Hz) was calculated as previous reported(Middleton et al., 2018). In Fig. 2h, only cells significantly phase locked (*P*<0.05, circular Rayleigh test) were shown, with 0º and 180º representing the peak and trough of SWRs separately. The mean phase-angle and mean resultant length (MRL) were calculated using CircStat. (circular statistic toolbox(Berens, 2009)).

### Single-unit and place cell properties

Units identified as pyramidal neurons were used for further analysis. The mean firing rate (FR) of each neuron was calculated as total number of spikes emitted by the neuron during Pre or Post sleep divided by session’s duration. The FR in SWRs was calculated as total number of spikes generated by the single unit divided by combined duration of all SWR events of the session. Participation of a single unit in SWRs was calculated as percent of SWR events the unit fired at least single spike in any given session. A burst was defined as two or more spikes occurring within a 10 ms time bin and all burst detection and analyses were performed as previously described(Bakkum et al., 2013). For place cell analyses, the firing rate maps were calculated by dividing the number of spikes falling into each 1cm × 1cm spatial bin by the total occupancy time of that bin and were subsequently smoothed with a 1 SD Gaussian kernel. This was repeated for both left and right direction on the linear track to generate a directional (left or right) firing rate curve. Mean firing rate was calculated by dividing the number of spikes occurring in periods where velocity exceeded 2 cm/s by that period’s duration and followed by averaging these values. Peak firing rates were defined as rate in the spatial bin having the maximal value. A place field was defined as a set of contiguous spatial bins surrounding the bin where the maximal firing rate was observed. In-field FR was defined as the total number of spikes emitted by the cell while the mouse was in the place field with the highest peak firing rate divided by the total time spent in this field, out-field FR was the number of spikes emitted by the cell in all spatial bins outside of the main field of that cell divided by the total time spent by the mouse outside of the main place field. Place field size was calculated by summing the number of spatial bins where a neuron’s firing rate exceed 20% of its peak firing rate. Spatial Information (bits per spike) was calculated as previously reported(Skaggs et al., 1993).

### Correlation analyses

To examine the coordination of spiking in CA1, CA2, and CA3 PCs during SWRs, spikes of individual units within a ±100 ms peri-SWRs window were pooled and z-scored firing rate curves (1 ms time bins) were constructed. We first calculated a correlation matrix of the normalized peri-SWRs firing rate curve of each neuron for Pre and Post session, using Pearson’s correlation coefficient (*r*). To validate the correlation matrix (Fig. 2c-d), both inter-region and intra-region pair-wise correlations were calculated. The intra-region correlation was defined as the Pearson’s correlation coefficient between two peri-SWRs firing rate curves of cells from the same region, while inter-region correlation was calculated for cells across different CA region (Extended Data Fig. 4a-b).

To quantify the temporal relation of ripples in CA1, CA2, and CA3, cross-correlations between SWR-peak (serving as the reference-event) and ripple power peaks of each region were calculated by generating a normalized SWR-triggered cross-correlation histograms (CCHs) using the same approach as previous described(Csicsvari et al., 2000; O’Neill et al., 2006). The confidence interval was determined by normal approximation as described previously(Frank et al., 2001; R, 1976), which allows for a simple quantification of confidence bounds without shuffling. We used a confidence interval of *P*<0.001 and only significantly correlated data were pooled to construct group averaged cross-correlations (Extended Data Fig. 2c).

To compare the structured pattern of spiking activity during the Pre- and Post-CNO session, a dendrogram was employed for hierarchical clustering of data from single Control and DREADD mouse (Extended Data Fig. 4c-d). Since CA1 and CA3 neurons discharged highly synchronously with the CA1 SWRs peak(Kay et al., 2016; Oliva et al., 2016a; Oliva et al., 2020), the peri-SWRs firing rate curves from CA1 and CA3 PCs were combined to generate a correlation matrix as described above using Pearson’s correlation coefficient. Then the pairwise Euclidian distance between each pair of observations was calculated (using MATLAB pdist function) and the linkage was decided by unweighted average distance (using MATLAB linkage function). We then clustered the values into separate groups and visualize them with dendrogram function with “ColorThreshold” of 70% of the maximum linkage. For representation of sorted correlation matrices, optimalleaforder function was applied and sorted the value accordingly.

### Cell assembly analyses

Cell assembly were detected using an unsupervised statistical framework based on a combination of principal component analysis (PCA) and independent component analysis (ICA) as previously described(Lopes-dos-Santos et al., 2013; Middleton et al., 2018; van de Ven et al., 2016). Spike trains for each CA1 pyramidal cells were binned into 25 ms time windows and normalized by z-score transform, followed by a construction of the correlation matrix for all neuron pairs. We then extracted assembly patterns in a two-step process. First, cell assemblies were identified by calculating the principal components of the resulting correlation matrix with eigenvalues above a threshold derived from an analytical probability function for uncorrelated data estimated by Marčenko-Pastur law. Next, independent component analysis (ICA) was applied to extract assembly patterns given by vector of weights representing the contribution of each cell to that assembly. Thus, cell assemblies on the linear track (LT1 or LT2) were identified as described above, and assembly reactivation was validated by tracking these assemblies over time in the subsequent sleep session as follows:

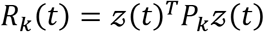

where *R*_*k*_(*t*) is the strength of assembly pattern *k* at time *t* and *z*(*t*) is the spike trains smoothed with a Gaussian kernel and subsequently z-score transformed as a function of time *t*. *P*_*k*_ is the projection matrix (outer product) of the corresponding independent component, and *T* is the transpose operator. To prevent the contribution of a single neuron’s high expression to assembly patters, the diagonal of *P*_*k*_ was set zero(Lopes-dos-Santos et al., 2013). By using this approach, only co-activation of multiple members of a cell assembly can contribute to an assembly pattern. Assembly activations were only included if *R* exceeded a threshold of five(van de Ven et al., 2016). Identified cell assemblies during Post-CNO session are as follows (N_Ctl-LT1_=82, N_DR-LT1_=36, N_Ctl-LT2_=72, N_DR-LT2_=30). Assembly strength on the linear track was calculated as the mean strength of all assembly events during mice traverse left and right laps. Assembly reactivation was measured by tracking the activity of the cell assemblies identified on linear track (LT1/LT2) during the following sleep session. Reactivation rate was defined as total number of activation events of individual assembly divided by duration. Reactivation strength was defined as the mean strength of all assembly events across session. Both reactivation rate and strength was assessed during SWR-periods.

To validate the time relation between assembly reactivation event and SWR event, we calculated the cross-correlations between assembly event-time and SWR-onset time (serving as reference-event) using a bin size of 5 ms. The confidence interval was determined by normal approximation as described above. Confidence interval was set to *P*<0.001 and only significant correlated data were pooled to generate group averaged cross-correlations (Fig. 4a-b). We also calculated a SWR active-index to measure how strongly a reactivation event was modulated by SWRs. The SWR active-index was defined as participation of an assembly event during SWRs at a time window of [T-10ms, T+10ms] divided by its participation at [T-50ms, T+50ms] serving as baseline, where T denotes SWR-onset time. Larger values represent reactivation events more temporal coincident with SWRs. When the baseline window was reduced to [T-30ms, T+30ms] or expanded to [T-100ms, T+100ms] consistent results were observed. For measurement of the temporal relationship between LT1 and LT2 assembly events during the Post-CNO session, the fraction of SWRs containing 3 categories of reactivation events (LT1-assembly only, LT2-assembly only, LT1 and LT2 assembly both) were quantified. Moreover, for every pair of LT1 and LT2 assembly events, a co-active strength was calculated as co-active rate of two assemblies during SWRs and then averaged across all SWR events. Pairwise assembly correlations between LT1 and LT2 were calculated using the Pearson’s correlation coefficient on 25-ms binned assembly trains. This was assessed inside and outside of SWRs as well. To quantify the event synchrony, we employed information theory and measured similarity and mutual dependence between LT1 and LT2 assemblies at the time domain. Mutual information, which quantity the joint information between two random variables, captures both linear and non-linear dependencies(Cover, 2006; Timme and Lapish, 2018). In this work, mutual information measures the synchrony between two assembly trains by binning assembly events into binary signals with a bin-size of 25 ms. Since there is no upper bound for mutual information, for comparison we normalized each value using the geometric mean as describe previously(Strehl, 2001), with 0 representing fully independence and 1 representing totally overlap between two assemblies. Joint entropy and conditional entropy were calculated as well to quantity the uncertainty between two variables(Cover, 2006; Timme and Lapish, 2018). Since both measure the randomness and certain state has zero entropy, its value is lower when two variables are more similar and dependent with each other.

### Replay analyses

We applied a Bayesian decoding approach to decode trajectory events as previous described(Davidson et al., 2009; Middleton et al., 2018; Silva et al., 2015). The ripple events detected in the Post-CNO session were further refined by excluding those whose duration was less than 80ms or events in which less than 10% of place cells participate. These ripples were subdivided into non-overlapping 20ms windows and firing rate vectors for all place cells were constructed within these bins. Bayesian decoding was then applied to generate a virtual position for each temporal window with the smoothed firing rates across space from the previous linear track (LT1/LT2) session serving as templates as follows:

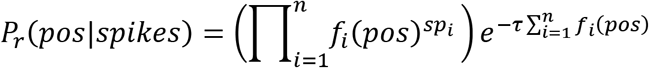

where *f*_*i*_(*pos*) is the firing rate in the template of the *i*^*th*^ place cell at position *pos, sp*_*i*_ is the number of spikes fired by the *i*^*th*^ place cell in the time-bin being decoded, *τ* denotes the duration of the time-bin (0.02s) and *n* is the total number of place cells. The posterior probabilities (*P*_*r*_) were then normalized to 1 using the following equation, where *P*_*n*_ is the total number of positions:

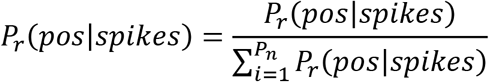

To estimate sequential structure, weighted correlation defined as Pearson’s correlation weighted by the decoded posterior probability was calculated first for each decoded event as previously described(Davidson et al., 2009; Middleton et al., 2018; Silva et al., 2015). Then, a Monte Carlo *P*-value was calculated to determine the significant replay sequences by circularly shifting firing rate curves at each time point for each place cell by a random distance. The Monte Carlo *P*-value was calculated as (*n*+1)/(*r*+1), where *r* is the total number of shuffles and *n* is the number of shuffles with a weighted correlation greater than or equal to the unshuffled value. Each candidate event was decoded twice using directional (left and right) smoothed firing curves, while its *P*-value equals to the smallest value yielding from left or right template. We performed 1000 times shuffle for each candidate and consider only events with a *P*-value below 0.05 to be significant replay events. Using this criteria, only 4%-7% of all analyzed candidates contained significant replay events (Ctl-LT1: 198/3710, 5.3%, Ctl-LT2: 263/3710, 7.1%, DREADD-LT1: 128/3271, 4%, DREADD-LT2: 154/3271, 4.7%; *P*_(LT1:Ctl>DR)_=0.005, *P*_(LT2:Ctl>DR)_<0.0001, chi-square test). To compare the replay quality across groups, two quantifications were used which include only significant replay events yielded above. A replay score which represents the estimated likelihood of an experienced trajectory by using a linear fitting algorithm was calculated as previously described(Davidson et al., 2009):

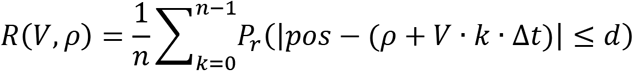

where *R*(*V*, *ρ*) represents the goodness-of-fit of the decoded events along the fitted line which is determined by slope *V* (velocity) and intercept *ρ* (starting location). For a candidate event consisting of *n* position estimates calculated at temporal bin size of Δ*t* (0.02s), the likelihood *R* that animal travels within distance *d* of a trajectory is determined by detecting the values of *V*_*max*_ and *ρ*_*max*_ that maximize *R*. *R*_*max*_ is the measure of the goodness-of-fit of the detected trajectory and the value of it denotes the replay score. The score ranges between 0 and 1 with a higher value representing a better line fit and higher replay quality.

The second approach to quantify replay content is sequence score (*rZ*) as previously described(Grosmark and Buzsaki, 2016). In brief, null (shuffled) firing rate curve for each place cell was constructed by circularly shifting its firing position by a random distance, and then used to decode event to generate a new shuffled weighted correlation. Thus, the full replay analysis (above) was performed 1000 times to generate 1000 *r*(*null*) values for each candidate. A final sequence score *rZ* was defined as:

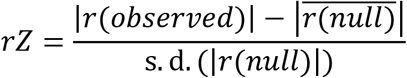

Additionally, a normalized max participation index was calculated to measure the degree of population discharge for each replay candidate. We calculated the maximum number of cells firing at least one spike in each time-bin (20ms) divided by total number of cells, resulting in a normalized max percentage of participation value for each replay event. The larger value indicates a stronger population synchrony during each time bin. Replay distance was defined as a set of peak posterior probability positions across time bins, the sharpness was defined as the amount of posterior probability concentrated near the peak as previous described(Silva et al., 2015).

### Statistical analysis

All statistical tests were performed using Matlab or SPSS. Unless otherwise noted, we used non-parametric two-sided Wilcoxon rank sum or signed rank test for group or pair-wise comparisons. For multiple comparisons, two-way ANOVA was conducted, and if necessary, followed by pair-wise post-hoc tests with Bonferroni’s correction. Specifically, besides interaction effect, post-hoc was conducted to validate which variables result in overall main effect. No statistical methods were used to predetermine sample sizes; instead, we opted to use group sizes similar to those in previously published studies(McHugh et al., 2007; Middleton and McHugh, 2016) to ensure variance was kept to a minimum between genotypes and cell quantities. Values are presented as mean±s.e.m. All boxplots represent the median (center marks) and the 25^th^-75^th^ percentiles, with whiskers extending to the most extreme data points without considering outliers, which were also included in statistical analyses. Unless noted, the level of statistical significance was set to 0.05 and *P* values were shown as follows: * *P* < 0.05; ** *P* < 0.01; *** *P* < 0.001.

## Acknowledgements

We thank Steven J. Middleton for providing input on the cell assembly and replay analyses and comments on the manuscript; M. Fujisawa and Y. Goto for daily assistance; the Advanced Manufacturing Support Team at RIKEN Center for Advanced Photonics for their assistance in microdrive production; all the members of the Laboratory for Circuit and Behavioral Physiology (CBP) for advice. This work was supported by Japan Society for the Promotion of Science (JSPS) Doctoral Course Fellowship (17J05573 to H.H.), a Grant-in-Aid for Scientific Research from MEXT (19H05646; T.J.M), a Grant-in-Aid for Scientific Research on Innovative Areas from MEXT (19H05233; T.J.M), and RIKEN CBS (T.J.M).

## Contributions

T.J.M conceived of the study and provided supervision along with K.O.. H.H. and T.J.M. analyzed data with input and assistance from R.B., E.T. and D.P.. R.B. collected the data, A.J.Y.H. produced all AAV vectors. H.H., and T.J.M wrote the paper with input from all authors.

## Corresponding authors

Correspondence to Thomas J McHugh.

---

**Extended Data Figure 1.**
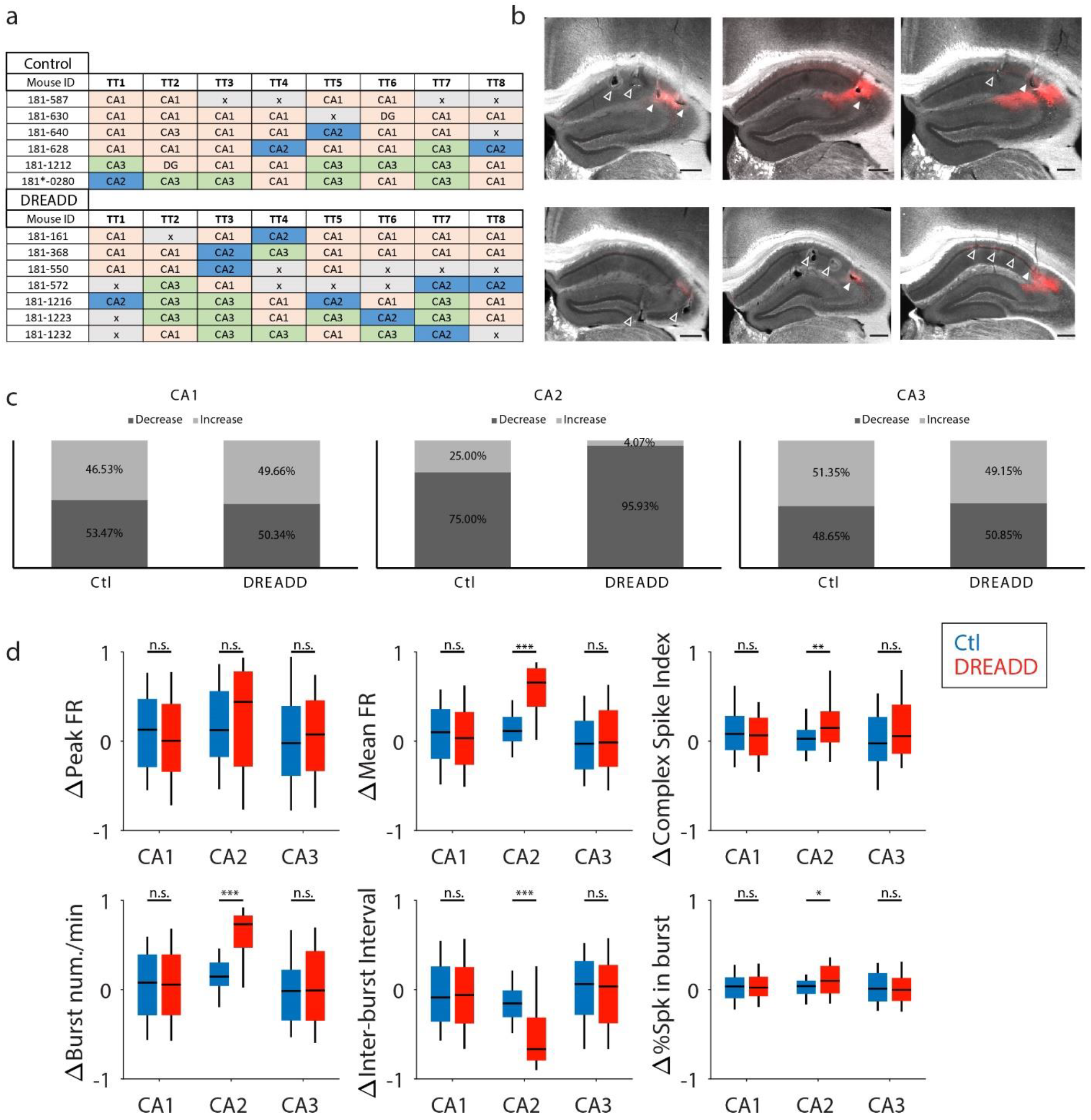
Alterations in hippocampal spiking following inhibition of CA2. **a.** Summary of all mice used for experiments with tetrode location coded with distinct colors. **b.** Bright field images of examples hippocampal slices indicating tetrode location and lesions in CA1, CA2, and CA3. CA2 was characterized by viral mCherry expression. Filled arrows indicate the recording site in CA2, while empty arrows illustrate CA1 or CA3 (Scale bars: 500 μm). **c.** Percent of cells that increased or decreased their firing rate (Pre vs, Post) in each CA region. Note a strong shutdown of firing was seen in CA2 of DREADD mice. **d.** Spike properties across all subfields. All the values were normalized as (Pre-Post)/(Pre+Post). Peak Firing rate (FR) showed no difference between groups in CA1 (*P*=0.126, *Z*=1.531), CA2 (*P*=0.103, *Z*=-1.632) and CA3 (*P*=0.741, *Z*=-0.331); Mean Firing rate (FR) showed a significant decrease only in CA2 (*P*=2.487×10^−10^, *Z*=-6.3278) in DREADD mice; Complex spike index (CSI) showed a significant decrease only in CA2 (*P*=0.005, *Z*=-2.788) in DREADD mice; Burst rate showed a significant decrease in CA2 (*P*=1.638×10^−11^, *Z*=-6.735) in DREADD mice; Inter-burst interval showed a significant increase in CA2 (*P*=8.936×10^−9^, *Z*=5.750) in DREADD mice; Percent of spike in burst showed a significant decrease in CA2 (*P*=0.029, *Z*=-2.1827) in DREADD mice. Statistical significance was tested using two-sided Wilcoxon rank-sum. All boxplots represent the median (center mark) and the 25th-75th percentiles, with whiskers extending to the most extreme data points without considering outliers, which were also included in statistical analyses; \**P*<0.05; \*\**P*<0.01; \*\*\**P*<0.001; n.s.= not significant

**Extended Data Figure 2.**
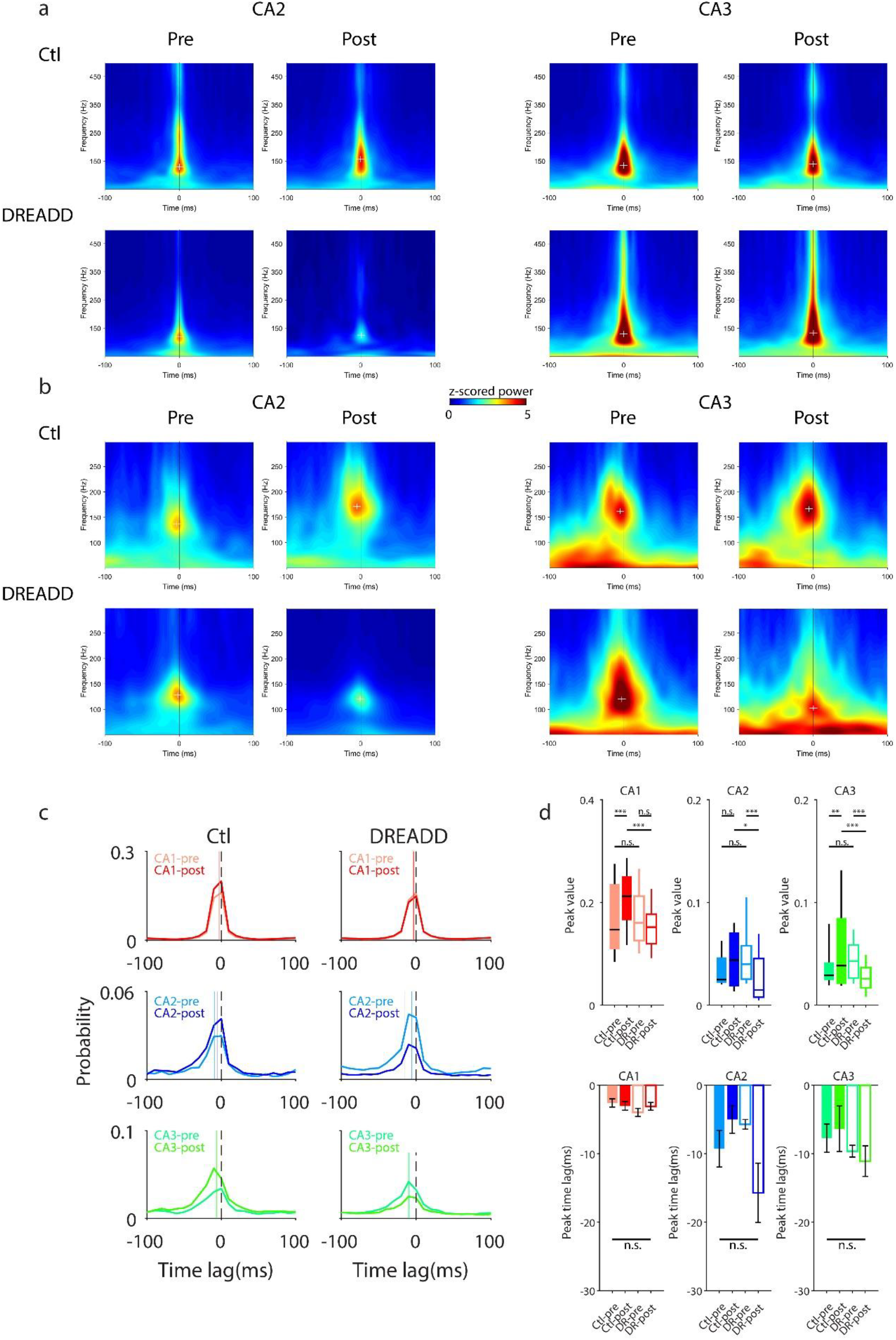
CA2 inhibition decreases the temporal coordination of ripples across the CA fields. **a.** Example of averaged SWR trigged spectrograms in CA2 and CA3 regions from Control (top row) and DREADD (bottom row) mice. The left panel (4 images) showed CA2 ripples using CA2 ripple peak as the temporal reference; the right panel (4 images) showed CA3 ripples using CA3 ripple peak as the temporal reference. Though the ripple power in each site was still aligned to the temporal reference, note a clear power decrease was seen in CA2 in DREADD mice. Black lines, ripple peak of each region; white cross, maximum ripple power for each region. **b.** Example of averaged SWR trigged spectrograms in CA2 and CA3 region using CA1 SWR peak as temporal reference for Control (top row) and DREADD (bottom row) mouse. The left panel (4 images) showed CA2 results; the right panel (4 images) showed CA3 results. Note a strong power decrease was observed both in CA2 and CA3 when aligned to the CA1 SWR-peak during Post-CNO session in DREADD mice. Black lines, CA1 SWRs peak; white cross, maximum ripple power for each region. **c.** Averaged population cross-correlation between SWR-peaks (serving as reference event) and ripple power peaks in each site for two groups. The dashed line denotes SWRs peaks, solid line indicates the ripple power peaks of each site. **d.** Quantification of cross-correlation in (**c**). The top panels show peak value and the bottom panels show absolute peak time lag (ms) for each region and group. For peak value comparisons in CA1, two-way ANOVA revealed a significant interaction (group × session) (*F*(1,242)=11.630, *P*=0.001), and post-hoc comparisons in each group revealed a significant increase in Controls (pre<post, *P*=0.001) but not in DREADD mice (pre vs. post, *P*=0.155), a significant decrease was also found in Post-CNO session (Ctl>DR, *P*<0.001) but not in Pre-CNO session (Ctl vs. DR, *P*=0.891); For peak value comparisons in CA2, a significant interaction (group × session) was also observed (*F*(1,122)=8.308, *P*=0.005), post-hoc comparisons revealed a significant decrease in DREADD (pre>post, *P*<0.001) but not in Controls (pre vs. post, *P*=0.226), a significant decrease was also found in Post-CNO session (Ctl>DR, *P*=0.020) but not in Pre-CNO session (Ctl vs. DR, *P*=0.088); For peak value comparisons in CA3, a significant interaction (*F*(1,150)=16.322, *P*<0.001) was found and post-hoc comparisons revealed a significant decrease in DREADD (pre>post, *P*=0.001) but a remarkable increase in Controls (pre<post, *P*=0.009). A significant difference was also found in Post-CNO session (Ctl>DR, *P*<0.001) but not in Pre-CNO session (Ctl vs. DR, *P*=0.322). There was no significant difference found in peak time lag across regions for the two groups. Tests were performed using two-way ANOVA with Bonferroni corrected post-hoc comparisons. All boxplots represent the median (center mark) and the 25th-75th percentiles, with whiskers extending to the most extreme data points without considering outliers, which were also included in statistical analyses; \**P*<0.05; \*\**P*<0.01; \*\*\**P*<0.001; n.s.= not significant

**Extended Data Figure 3.**
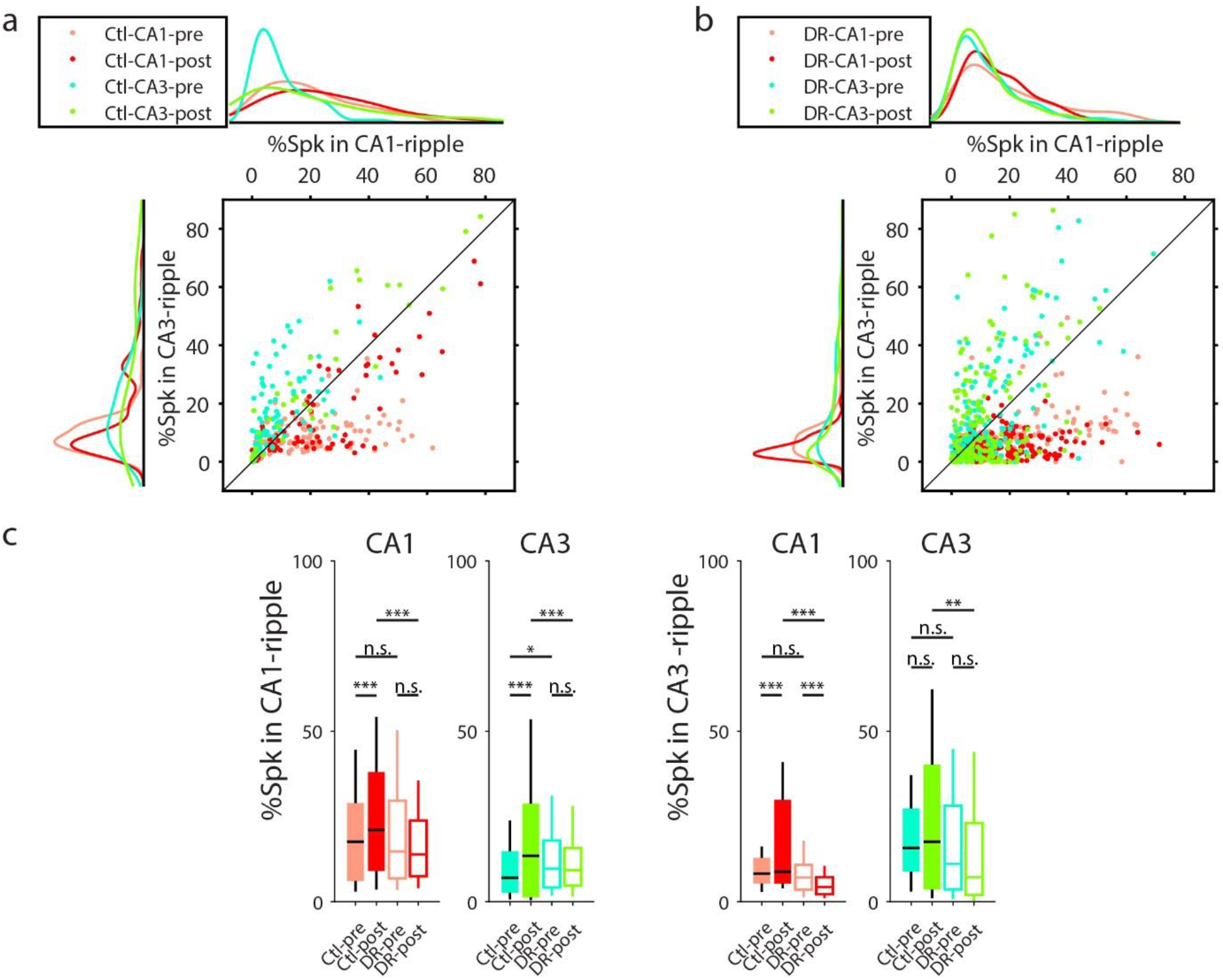
Disrupted CA1-CA3 spike interaction during SWRs in CA2 DREADD mice. **a.** Percent of CA1 and CA3 spikes inside CA1 and CA3 ripples during the Pre- and Post-CNO session in Control mice. Compared to the Pre-CNO session, a greater number of CA1 and CA3 spikes were involved in CA1 and CA3 ripples during Post-CNO session, following spatial exploration on the track.. **b.** Percent of CA1 and CA3 spikes inside CA1 and CA3 ripples during Pre- and Post-CNO session in DREADD mice. No obvious change of CA1 and CA3 spikes involved in CA1 and CA3 ripples comparing Pre- and Post-CNO session. For **a.** and **b.**, only cells active both in CA1 and CA3 ripples were included. **c.** Quantification of **a**. and **b.** shown above. The left panel showed the percent of CA1 and CA3 spikes occurring inside CA1 ripples, the right panel showed the percent of CA1 and CA3 spikes inside CA3 ripples. For % of CA1 spikes inside CA1 ripples, two-way ANOVA revealed a significant interaction (group × session) (*F*(1,506)=9.141, *P*=0.003) and post-hoc comparisons revealed a significant increase in Controls (pre<post, *P*=0.016) but not in DREADD mice (pre vs. post, *P*=0.071), a significant difference was also found in Post-CNO session (Ctl>DR, *P*<0.001) but not in Pre-CNO session (Ctl vs. DR, *P*=0.885). For % of CA3 spikes inside CA1 ripples, two-way ANOVA revealed a significant interaction (group × session) (*F*(1,434)=18.415, *P*<0.001), and post-hoc comparisons revealed a significant increase in Controls (pre<post, *P*<0.001) but not in DREADD mice (pre vs. post, *P*=0.306), a strong effect was found also in Post-CNO session (Ctl>DR, *P*<0.001) with a minor difference in Pre-CNO session (Ctl<DR, *P*=0.022). For % of CA1 spikes inside CA3 ripples, two-way ANOVA revealed a significant interaction (*F*(1,506)=50.220, *P*<0.001); post-hoc comparisons revealed a significant effect in Controls (pre<post, *P*<0.001) but opposite effect was found in DREADD mice (pre>post, P<0.001), a significant difference was found also in Post-CNO session (Ctl>DR, *P*<0.001) but not in Pre-CNO session (Ctl vs. DR, *P*=0.519). For % of CA3 spikes inside CA3 ripples, two-way ANOVA revealed a significant interaction (*F*(1,434)=5.341, *P*=0.021), and post-hoc comparisons revealed a significant difference in the Post-CNO session (Ctl>DR, *P*=0.002) but not in Pre-CNO session (Ctl vs. DR, *P*=0.823). All boxplots represent the median (center mark) and the 25th-75th percentiles, with whiskers extending to the most extreme data points without considering outliers, which were also included in statistical analyses; \**P*<0.05; \*\**P*<0.01; \*\*\**P*<0.001; n.s.= not significant

**Extended Data Figure 4.**
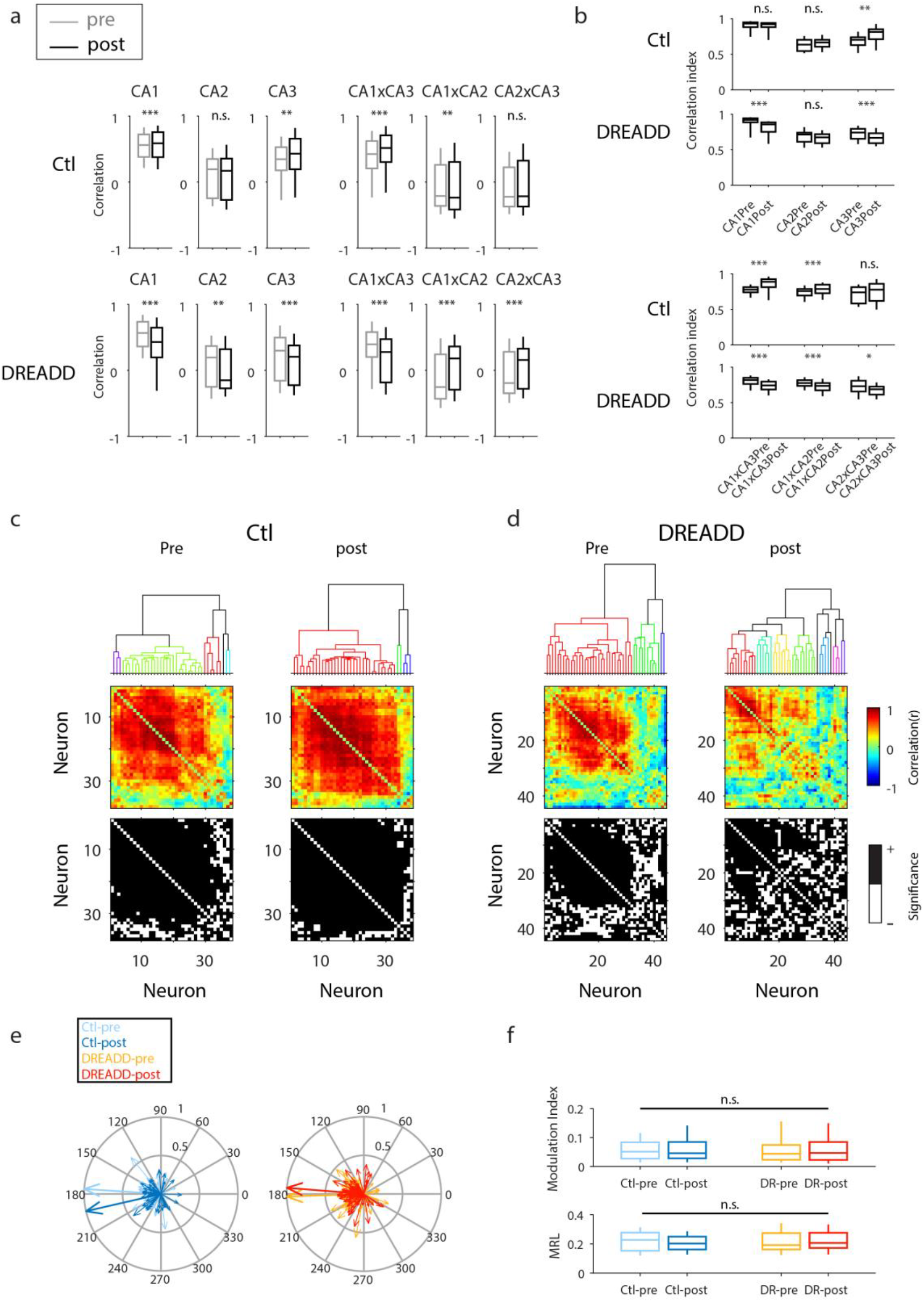
Diminished SWR-associated spike correlation in CA2 DREADD mice. **a.** Intra-regional and inter-regional correlation for peri-SWR mean firing rate curves for neurons in CA1, CA2, and CA3 in Pre-CNO (grey) and Post-CNO (black) session for Control (top row) and DREADD mice (bottom row). For intra-regional correlation, a significant increase of correlation was found in CA1 and CA3 (CA1: pre<post, *P*=6.947×10^−6^, *Z*=4.495; CA3: pre<post, *P*=0.001, *Z*=3.269), not for CA2 (pre vs. post, *P*=0.470, *Z*=-0.722) in Control mice; whereas in DREADD mice, a significant decrease of correlation was found in all CA sites (CA1: pre>post, *P*<0.001, *Z*=-40.596; CA2: pre>post, *P*=0.003, *Z*=-2.929; CA3: pre>post, *P*=2.909×10^−35^, *Z*=-12.391). For inter-regional correlation, Control mice displayed a significant increase of correlation in CA1xCA3 (pre<post, *P*=2.055×10^−27^, *Z*=10.847), a decrease of correlation in CA1xCA2 (pre>post, *P*=0.004, *Z*=-2.906), and comparable in CA2xCA3 (pre vs. post, *P*=0.100, *Z*=1.646). However, DREADD mice showed a significant decrease of correlation in CA1xCA3 (pre>post, *P*=3.460×10^−226^, *Z*=-32.108), an increase of correlation in CA1xCA2 (pre<post, *P*=3.693×10^−169^, *Z*=27.723) and CA2xCA3 (pre<post, *P*=1.390×10^−23^, *Z*=10.009).Two-sided Wilcoxon rank-sum test. **b.** Quantification of the correlation matrix in **Fig. 2c-d**. The correlation index was calculated as (#significant correlated cells)/(#total cells) in Pre- and Post-CNO session separately. For the intra-region comparisons in Control mice, two-sided Wilcoxon rank-sum revealed a significant increase in CA3 (pre<post, *P*=0.0026, *Z*=-3.1053), while CA1 (pre vs. post, *P*=0.114, *Z*=2.831) and CA2 (pre vs. post, *P*=0.346, *Z*=-0.942) were similarly unchanged. However, in DREADD mice, a significant decrease in CA1 (pre>post, *P*=1.121×10^−29^, *Z*=11.314) and CA3 (pre>post, *P*=1.009×10^−4^, *Z*=3.888) was observed, while values remained similar in CA2 (pre vs. post, *P*=0.184, *Z*=1.329). For the inter-region comparisons, Control mice displayed a significant increase in CA1xCA3 (pre<post, *P*=1.239×10^−14^, *Z*=-7.712) and CA1xCA2 (pre<post, *P*=4.119×10^−5^, *Z*=-4.101) correlations, while correlations remained similar in CA2xCA3 (pre vs. post, *P*=0.224, *Z*=-1.217). In contrast, CA2 DREADD mice displayed a significant decrease in correlation in CA1xCA3 (pre>post, *P*=1.424×10^−19^, *Z*=9.050), CA1xCA2 (pre>post, *P*=2.507×10^−9^, *Z*=5.961), and CA2xCA3 (pre>post, *P*=0.018, *Z*=2.368) pairs. **c.** Example of hierarchical clustering of correlation matrix generated by the peri-SWRs firing rate curves of CA1 and CA3 pyramidal neurons from one Control mouse during the Pre- and Post-CNO session. Top, dendrogram with different colors coded for different clusters. Middle, correlation matrix sorted by matching the order in the dendrogram. Bottom, significant value of correlation matrix shown in the Middle, with black representing significant correlation value while white representing not significant. Note that following exploration of the novel LT2 track, a more structured and clustered pattern was observed in the controls during the Post-CNO compared to the Pre-CNO sessions, and a clear increase number of significant pairs were observed also during the Post-CNO session. **d.** Example of hierarchical clustering of correlation matrix generated by the peri-SWRs firing rate curves of CA1 and CA3 pyramidal neurons from one DREADD mouse during the Pre- and Post-CNO session. Note that the structured pattern broke into more clusters in Post-CNO compared to the Pre-CNO session, accompanied with more non-significant cell-pairs. **e.** Polar plots indicate the mean phase angle at which all CA1 PCs fires during SWRs in Pre- and Post-CNO session. Each neuron was shown as a thin arrow, the length illustrates the mean resultant length (MRL). 0 and 180 degree indicate the peak and trough of ripple oscillation separately. Group means were shown as thick arrows but indicate phase only (preferred phase: Ctl-pre: 174.650±8.230°, Ctl-post: 195.830±9.514°, DREADD-pre: 179.953±6.229°, DREADD-post: 180.725±6.095°). **f.** Quantification of ripple phase analysis. Control and DREAD mice showed similar in preferred phase (*P*=0.429, *F*=0.925, circular ANOVA), mean resultant length (MRL: *P*=0.608, *F*=0.611, ANOVA) and Modulation Index (MI: *P*=0.929, *F*=0.1506, ANOVA). All boxplots represent the median (center mark) and the 25th-75th percentiles, with whiskers extending to the most extreme data points without considering outliers, which were also included in statistical analyses; \**P*<0.05; \*\**P*<0.01; \*\*\**P*<0.001; n.s.= not significant

**Extended Data Figure 5.**
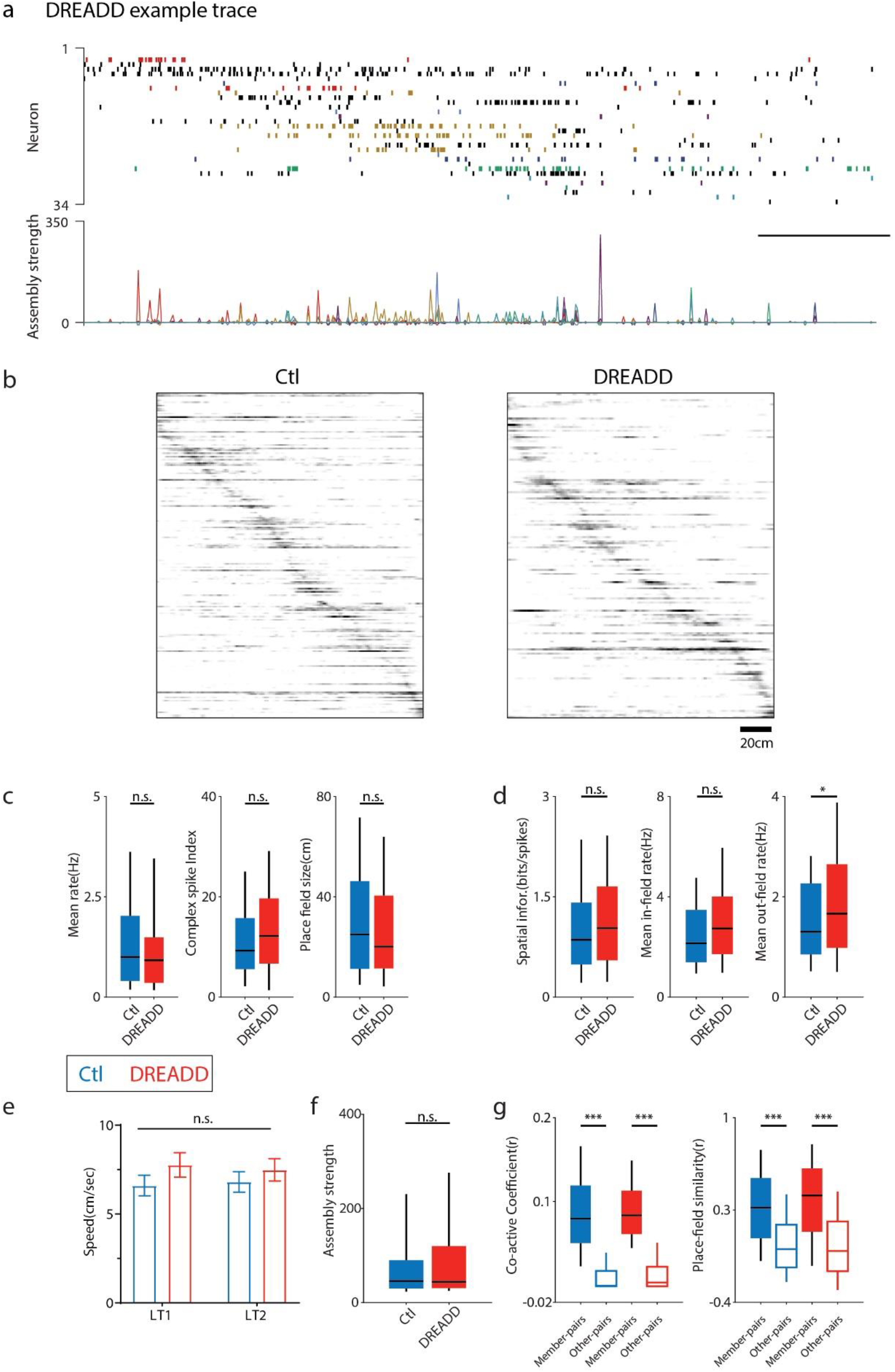
Spatial rate coding was comparable for Control and CA2 DREADD mice. **a.** Example of CA1 neuronal activity during a single lap running on the LT2 for the DREADD mouse. The raster plot shows the firing activity during this lap of 34 simultaneously recorded pyramidal cells with different colors representing neurons identified as each single cell assembly (7 cell assembly overall). Spikes in black are shown for completeness but do not belong to any significantly detected cell assemblies. The expression strength of each cell assembly was shown on the bottom. The horizontal scale bar indicates 2 s. **b.** Place field rate maps sorted by their peak firing position on LT2 for Control mice (left panel) and DREADD mice (right panel). **c, d.** Spatial rate coding during track exploring (LT2) in CA1 is unaltered despite the inhibition of CA2. Place cells of Control and DREADD mice showed comparable firing properties in mean FR (*P*=0.395, *Z*=0.850), CSI (*P*=0.071, *Z*=-1.809), place field size (*P*=0.140, *Z*=1.475), spatial information (*P*=0.233, *Z*=-1.192), and FR within place field (*P*=0.072, *Z*=-1.802), while there was a significant increase in FR outside the place field in the CA2 DREADD mice (*P*=0.035, *Z*=-2.107). All tests are performed using two-sided Wilcoxon rank-sum. **e.** No differences in running speed were observed between control and DREADD mice on the track (*P*>0.05, two-way ANOVA) **f.** Mean assembly strength during exploration on LT2 was comparable between groups (Ctl: 93.346±13.105, DREADD: 111.639±30.370, *P*=0.885, Z=-0.145, two-sided Wilcoxon rank-sum test) **g.** (Left) Place-cell pairs on LT2 belonging to the same assembly (member-pairs) showed a higher co-firing coefficient (Pearson’s r) than pairs which do not belong to part of any assembly (other-pairs) in both control and DREADD mice (two-way ANOVA with Bonferroni’s multiple-comparisons: group (Ctl vs. DR): *F*(1,3198)=2.006, *P*=0.157; type (member vs. other): *F*(1,3198)=1459.167, *P*<0.001; interaction: *F*(1,3198)=0.405, *P*=0.525; post-hoc: Ctl: *P*<0.001 (member>other), DR: *P*<0.001 (member>other)). (Right) Place field similarities, calculated as the Pearson’s correlation between cell pair rate maps for member or other pairs, was also higher for member-pairs than other-pairs for both groups (two-way ANOVA with Bonferroni’s multiple-comparisons: group (Ctl vs. DR): *F*(1,3198)=1.247, *P*=0.264; type (member vs. other): *F*(1,3198)=215.549, *P*<0.001; interaction: *F*(1,3198)=2.688, *P*=0.101; post-hoc: Ctl: *P*<0.001 (member>other), DR: *P*<0.001 (member>other)). All boxplots represent the median (center mark) and the 25th-75th percentiles, with whiskers extending to the most extreme data points without considering outliers, which were also included in statistical analyses; bar graphs represent mean ± s.e.m.; \**P*<0.05; \*\**P*<0.01; \*\*\**P*<0.001; n.s.= not significant

**Extended Data Figure 6.**
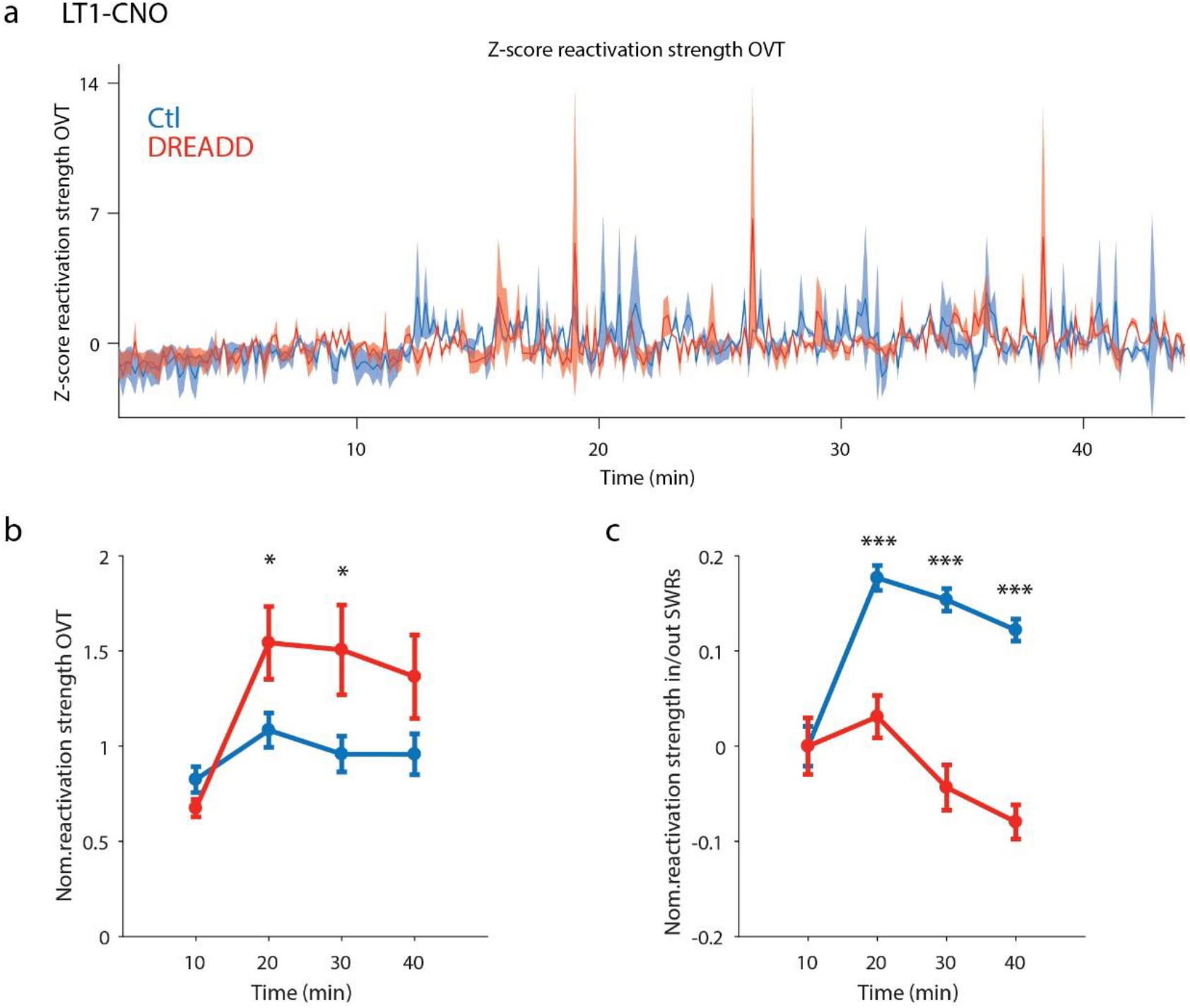
Decreased accuracy of SWR-associated assembly reactivation during the CNO session in DREADD mice. **a.** Averaged reactivation strength (mean ± sem) changing overtime (OVT) for the two groups. The assemblies in the CNO session were identified from the LT1 session. CNO was administrated at time point 0. A comparable reactivation was observed during the first 10 min in both groups, however during the second 10 min time-bin increased reactivation strength could be seen in the DREADD mice. **b.** Averaged normalized reactivation strength at 4 time-bins throughout the session. The value was normalized to the baseline (first 10 mins). Control and DREADD mice were comparable in 10 min bin (*P*=0.133, *Z*=1.504) and 40 min bin (*P*=0.260, *Z*=-1.127), yet DREADD displayed higher reactivation than Control mice in 20 min bin (*P*=0.024, *Z*= −2.265) and 30 min bin (*P*=0.023, *Z*= −2.281), indicating a reliable CNO dependent change. Tests were performed using two-sided Wilcoxon rank-sum at each time point. **c.** Averaged normalized reactivation strength at 4 time-bins during in/out SWR-period. The strength ratio of each assembly was firstly calculated as Strength-within-SWR/(Strength-within-SWR + Strength-out-SWR), then normalized to the baseline (first 10 mins). Control and DREADD mice were similar at 10 min bin (*P*=0.844, *Z*=-0.197), whereas a remarkable decrease was found in 20 min bin (*P*=8.398×10^−7^, *Z*=4.926), 30 min bin (*P*=6.866×10^−11^, *Z*=6.524) and 40 min bin (*P*=1.481×10^−13^, *Z*=7.389) in DREADD mice, indicating an impairment of SWR-associated assembly reactivation following CNO administration. Tests were performed using two-sided Wilcoxon rank-sum at each time point. All data are represented as mean ± s.e.m.; \**P*<0.05; \*\**P*<0.01; \*\*\**P*<0.001; n.s.= not significant

**Extended Data Figure 7.**
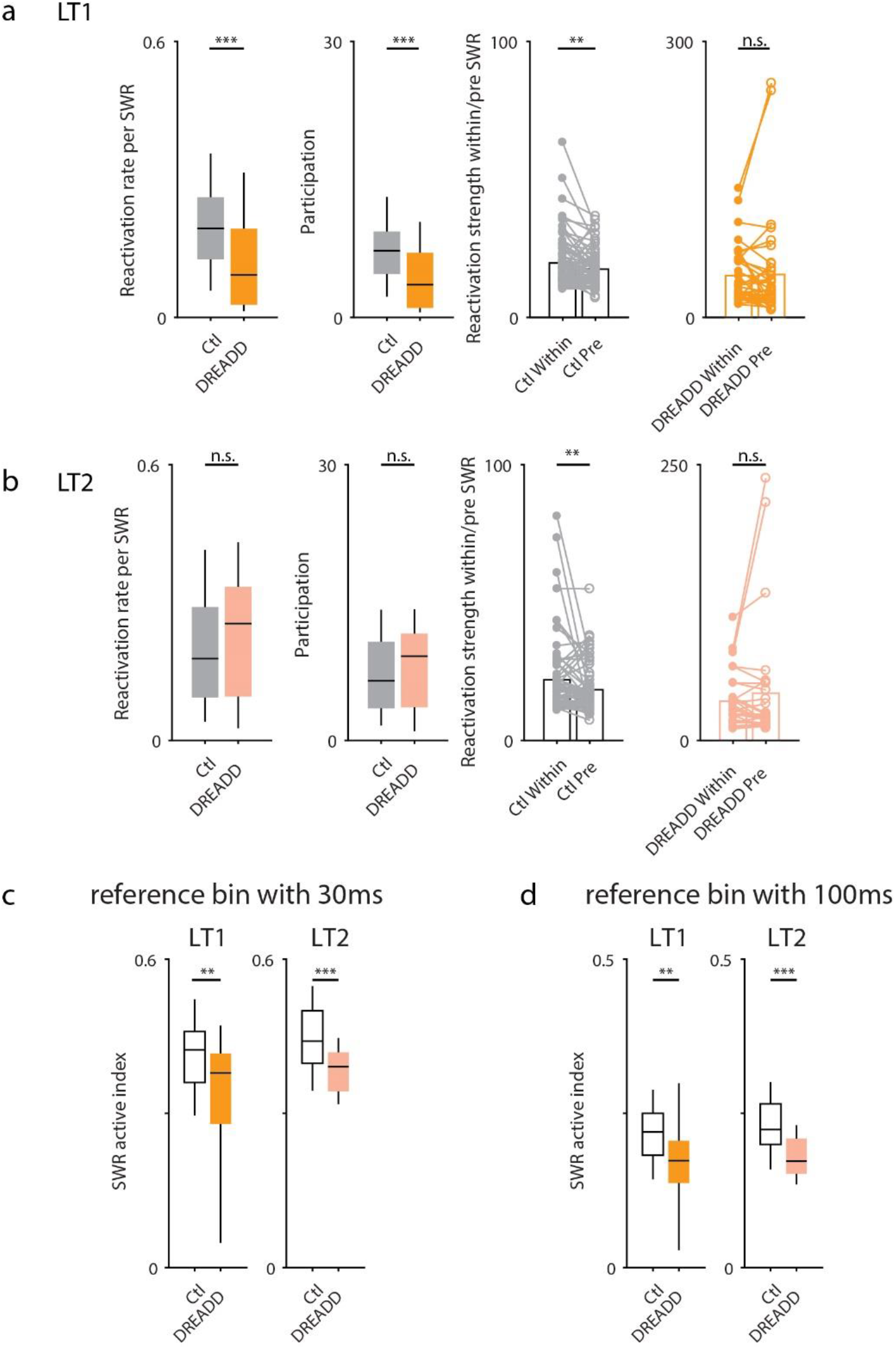
Altered assembly reactivation properties in the CA2 DREADD mice. **a.** LT1 assembly properties during the Post-CNO session. DREADD mice displayed a significant decrease in reactivation rate per SWR (Ctl>DR: *P*=3.928×10^−4^, *Z*=3.545, Wilcoxon rank-sum) and participation of SWRs (Ctl>DR: *P*=1.991×10^−4^, *Z*=3.720, Wilcoxon rank-sum) compared to Controls. Further, a significant difference was found between reactivation strength within-SWR and pre-SWR (200ms pre-window) in Control mice (within-SWR>pre-SWR: *P*=0.001, *Z*=3.266, Wilcoxon sign-rank), whereas the values were comparable in DREADD mice (within-SWR vs. pre-SWR: *P*=0.322, *Z*=0.990, Wilcoxon sign-rank). LT2 assembly properties during the Post-CNO session. DREADD mice exhibited comparable reactivation rate per SWR (Ctl vs. DR: *P*=0.216, *Z*=-1.238, Wilcoxon rank-sum) and participation of SWRs (Ctl vs. DR: *P*=0.285, *Z*=-1.069, Wilcoxon rank-sum) to Controls. A significant difference was found between reactivation strength within-SWR and pre-SWR (200ms pre-window) in Control mice (within-SWR>pre-SWR: *P*=0.002, *Z*=3.086, Wilcoxon sign-rank), whereas values were similar in DREADD mice (within-SWR vs. pre-SWR: *P*=0.371, *Z*=0.895, Wilcoxon sign-rank). **c.** SWR active-index taking the reference bin as 30 ms. Significant lower value was observed comparing the DREADD with the Control mice both for LT1 (Ctl>DR: *P*=0.003, *Z*=2.966) and LT2 (Ctl>DR: *P*=7.405×10^−4^, *Z*=3.374), which is consistent with the result of **Fig. 4b**. Two-sided Wilcoxon rank-sum. **d.** SWR active-index taking the reference bin as 100 ms. Significant lower value was observed comparing the DREADD with the Ctl mice both for LT1 (Ctl>DR: *P*=0.002, *Z*=3.169) and LT2 (Ctl>DR: *P*=1.057×10^−4^, *Z*=3.877), which is also consistent with the result of **Fig. 4b** and **Extended Data Fig. 7c**. Two-sided Wilcoxon rank-sum. All boxplots represent the median (center mark) and the 25th-75th percentiles, with whiskers extending to the most extreme data points without considering outliers, which were also included in statistical analyses; individual data points each represent properties of a single cell assembly; \**P*<0.05; \*\**P*<0.01; \*\*\**P*<0.001; n.s.= not significant

**Extended Data Figure 8.**
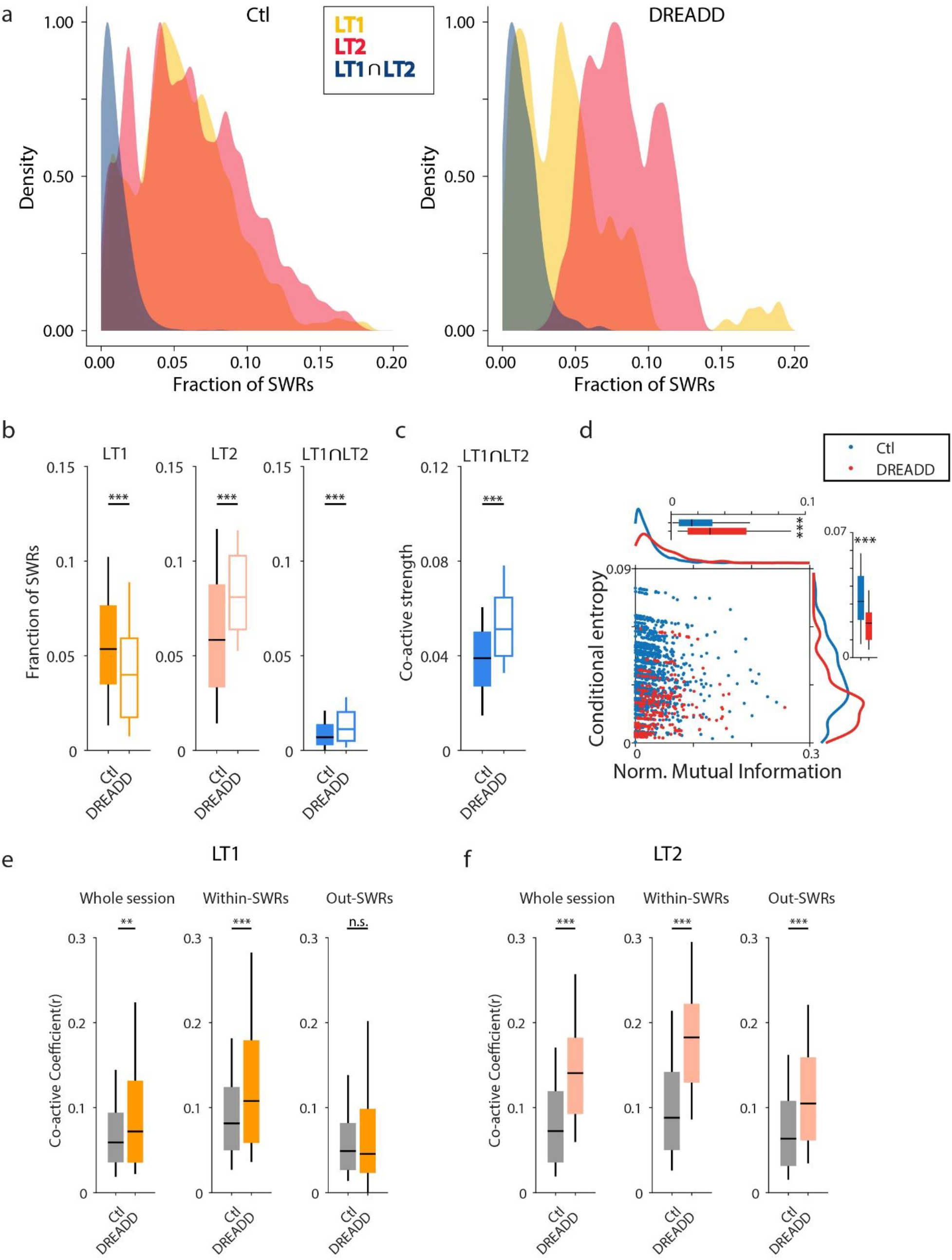
Decreased specificity of assembly reactivation in CA2 DREADD mice. **a.** Density plots of the fraction of SWRs for 3 categories (containing LT1-assembly only, containing LT2-assembly only, and containing both LT1 and LT2 assembly). Note the increase of SWRs containing both LT1 and LT2 assemblies in the DREADD mice. **b.** Quantifications for fraction of SWRs above. DREADD mice displayed a significantly lower fraction of SWRs containing LT1 assembly only (Ctl>DR: *P*=3.750×10^−12^, *Z*=6.946), a higher fraction of SWRs containing LT2 assembly only (Ctl<DR: *P*=1.27×10^−24^, *Z*=-10.243), and a higher fraction of SWRs containing both LT1 and LT2 (Ctl<DR: *P*=1.154×10^−12^, *Z*=-7.111), indicating a reduction in the precision and specificity of assembly reactivation in DREADD mice. Two-sided Wilcoxon rank-sum. **c.** Co-active strength representing the degree of co-activity for two assemblies, revealed a significant increase in DREADD mice compared to Controls (Ctl<DR: *P*=6.371×10^−5^, *Z*=-3.999). Two-sided Wilcoxon rank-sum. **d.** Scatter plot showing the coordination between LT1 and LT2 assemblies for two groups throughout the Post-CNO session. DREADD mice showed a significant increase in normalized mutual information (Ctl<DR: *P*=8.393×10^−18^, *Z*=-8.594) but a decrease in conditional entropy (Ctl>DR: *P*=1.342×10^−36^, *Z*=12.636), reflecting a reduction in track specificity and independency of the two representations. Each point represents an assembly pair. Two-sided Wilcoxon rank-sum. **e.** Assembly pair-wise correlation in Post-CNO session for LT1 assemblies. DREADD mice displayed significantly higher correlation across the session (Ctl<DR: *P*=0.006, *Z*=-2.759) and within-SWRs period (Ctl<DR: *P*=3.010×10^−6^, *Z*=-4.670) than Control mice, while comparable values were observed during the out-SWRs period (Ctl vs. DR: *P*=0.803, *Z*=0.250), indicating higher synchrony for LT1 assemblies in DREADD mice. Two-sided Wilcoxon rank-sum. **f.** Assembly pair-wise correlation in Post-CNO session for LT2 assemblies. DREADD mice displayed significantly higher correlation than Control mice across the session (Ctl<DR: *P*=3.490×10^−18^, *Z*=-8.694), within-SWRs period (Ctl<DR: *P*=2.114×10^−23^, *Z*=-9.968) and out-SWRs period (Ctl<DR: *P*=4.760×10^−9^, *Z*=-5.855), indicating higher synchrony for LT2 assemblies in DREADD mice. Two-sided Wilcoxon rank-sum. All boxplots represent the median (center mark) and the 25th-75th percentiles, with whiskers extending to the most extreme data points without considering outliers, which were also included in statistical analyses; \**P*<0.05; \*\**P*<0.01; \*\*\**P*<0.001; n.s.= not significant

**Extended Data Figure 9.**
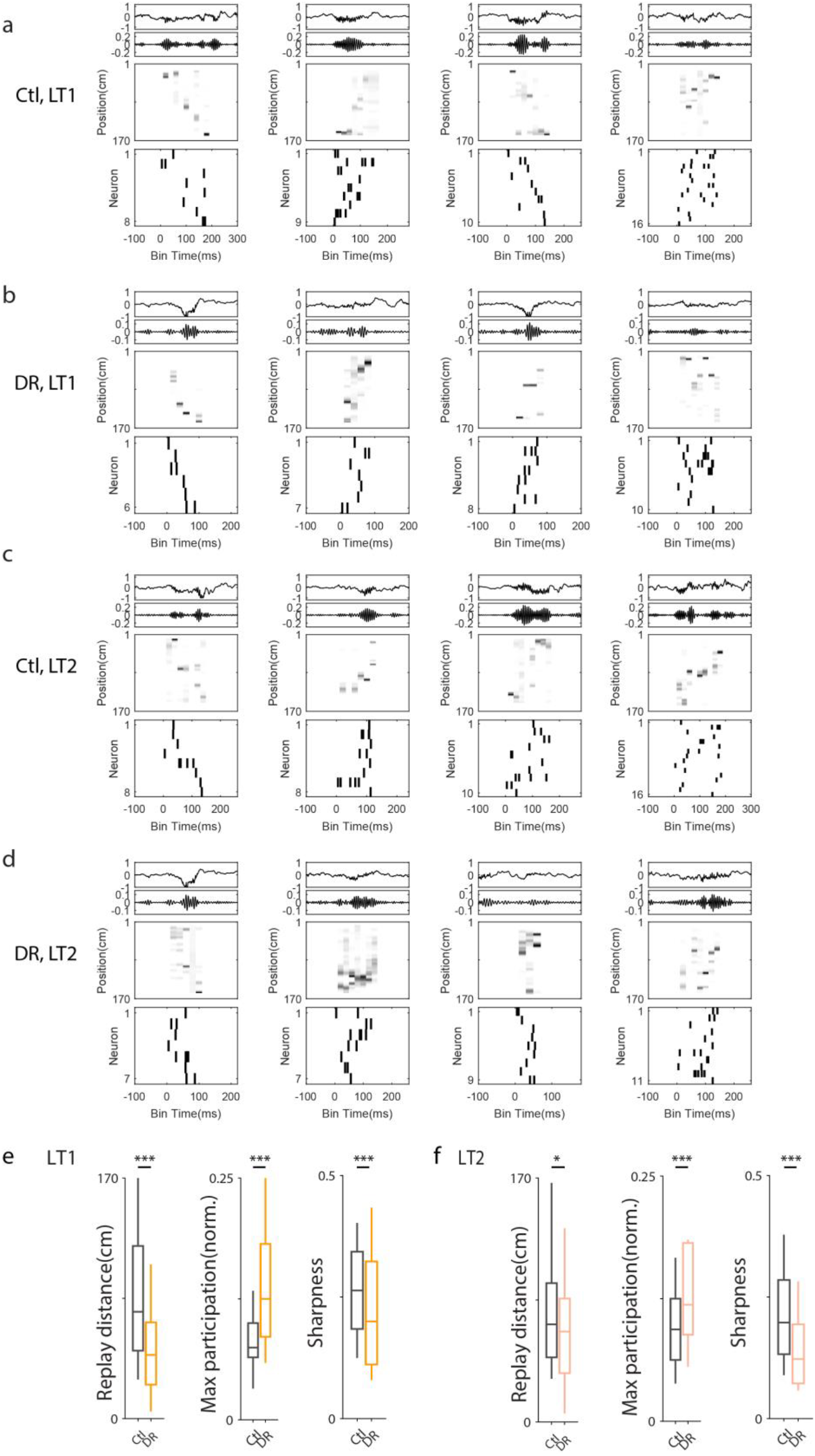
Poorer replay content and stronger population synchrony in DREADD mice. **a-d**. Examples of replay candidate events during SWRs in Post-CNO session for Control and DREADD mice. For each example, the following is shown from top to bottom: wideband LFP; LFP filtered in the SWR band; reconstructed replay structure; associated spike raster plot sorted based on firing location on each track. **e.** Quantification of LT1 sequences. DREADD mice showed a significant reduction in replay distance (Ctl>DR: *P*=7.418×10^−11^, *Z*=6.512) and sharpness (Ctl>DR: *P*=5.003×10^−4^, *Z*=3.481), with a significantly larger proportion of cells participating in each time-bin (Ctl<DR: *P*=4.992×10^−13^, *Z*=-7.226) compared to Control mice, indicating truncated trajectories accompanied with higher population synchronization at each time bin. **f.** Quantification of LT2 sequences. DREADD mice showed a significant reduction in replay distance (Ctl>DR: *P*=0.023, *Z*=2.280) and sharpness (Ctl>DR: *P*=9.345×10^−13^, *Z*=7.140), with a significant increase in the proportion of cells participating in each time-bin (Ctl<DR: *P*=8.686×10^−7^, *Z*=-4.919) compared to Control mice, consistent with LT1 results (**e**), also indicating the shortened replay span accompanied with higher population synchronization at each time bin leads to poorer replay content in LT2 sequence in DREADD mice. All boxplots represent the median (center mark) and the 25th-75th percentiles, with whiskers extending to the most extreme data points without considering outliers, which were also included in statistical analyses; \**P*<0.05; \*\*\**P*<0.001

