## Supplemental Table 1 for "CA2 inhibition reduces the precision of hippocampal assembly reactivation"

| Part | Content | Test | Mark | Statistic details | CA1(mean±sem) | CA2(mean±sem) | CA3(mean±sem) |
|---|---|---|---|---|---|---|---|
| c | Mean FR | Two-way ANOVA with Bonferroni's multiple comparison test | n.s. | (CA1) group: F(1,902)=1.176, P=0.279; session: F(1,902)=3.728, P=0.054; group x session: F(1,902)=1.680, P=0.195 | Ctl-pre: 0.789±0.093 Hz<br>Ctl-post: 0.578±0.058 Hz<br>DR-pre: 0.633±0.056 Hz<br>DR-post: 0.592±0.048 Hz | Ctl-pre: 0.651±0.113 Hz<br>Ctl-post: 0.489±0.085 Hz<br>DR-pre: 0.658±0.072 Hz<br>DR-post: 0.361±0.135 Hz | Ctl-pre: 0.429±0.105 Hz<br>Ctl-post: 0.375±0.082 Hz<br>DR-pre: 0.417±0.054 Hz<br>DR-post: 0.376±0.054 Hz |
| | | | sig | (CA2) group: F(1,234)=0.287, P=0.593; session: F(1,234)=4.094, P=0.044; group x session: F(1,234)=0.349, P=0.555. Post-hoc: P=0.368 (Ctl: pre vs. post); P=0.033 (DR: pre vs.post) | | | |
| | | | n.s. | (CA3) group: F(1,394)=0.006, P=0.941; session: F(1,394)=0.387, P=0.534; group x session: F(1,394)=0.006, P=0.938 | | | |
| e | FR in SWR | Two-way ANOVA with Bonferroni's multiple comparison test | sig | (CA1) group: F(1,1051)=0.296, P=0.587; session: F(1,1051)=7.679, P=0.006; group x session: F(1,1051)=0.021, P=0.884. Post-hoc: P=0.072 (Ctl: pre vs. post); P=0.025 (DR: pre vs.post) | Ctl-pre: 1.874±0.154 Hz<br>Ctl-post: 1.522±0.119 Hz<br>DR-pre: 1.771±0.101 Hz<br>DR-post: 1.474±0.096 Hz | Ctl-pre: 0.708±0.230 Hz<br>Ctl-post: 0.462±0.162 Hz<br>DR-pre: 0.646±0.113 Hz<br>DR-post: 0.281±0.044 Hz | Ctl-pre: 0.891±0.139 Hz<br>Ctl-post: 0.999±0.122 Hz<br>DR-pre: 0.721±0.099 Hz<br>DR-post: 0.409±0.052 Hz |
| | | | sig | (CA2) group: F(1,178)=0.828, P=0.364; session: F(1,178)=5.223, P=0.023; group x session: F(1,178)=0.198, P=0.657. Post-hoc: P=0.247 (Ctl: pre vs. post); P=0.027 (DR: pre vs.post) | | | |
| | | | sig | (CA3) group: F(1,426)=11.274, P=0.001; session: F(1,426)=0.736, P=0.391; group x session: F(1,426)=3.355, P=0.068. Post-hoc: P=0.213 (pre: Ctl vs. DR); P=0.001 (post: Ctl vs. DR) | | | |
| f | Participation(%) | Two-way ANOVA with Bonferroni's multiple comparison test | sig | (CA1) group: F(1,1051)=3.666, P=0.056; session: F(1,1051)=4.709, P=0.030; group x session: F(1,1051)=0.079, P=0.779. Post-hoc: P=0.244 (Ctl: pre vs. post); P=0.037 (DR: pre vs.post) | Ctl-pre: 21.168±1.192<br>Ctl-post: 19.367±1.144<br>DR-pre: 19.387±0.779<br>DR-post: 17.274±0.731 | Ctl-pre: 11.078±1.577<br>Ctl-post: 7.828±1.466<br>DR-pre: 12.357±1.062<br>DR-post: 5.577±0.868 | Ctl-pre: 10.684±1.008<br>Ctl-post: 12.666±1.754<br>DR-pre: 8.803±0.809<br>DR-post: 5.490±0.490 |
| | | | sig | (CA2) group: F(1,209)=0.151, P=0.698; session: F(1,209)=16.099, P<0.001; group x session: F(1,209)=1.993, P=0.160. Post-hoc: P=0.103 (Ctl: pre vs. post); P<0.001 (DR: pre vs.post) | | | |
| | | | sig | (CA3) group: F(1,438)=21.825, P<0.001; session: F(1,438)=0.471, P=0.493; group x session: F(1,438)=7.456, P=0.007. Post-hoc: P=0.237 (Ctl: pre vs. post); P=0.001 (DR: pre vs. post); P=0.114 (pre: Ctl vs. DR); P<0.001 (post: Ctl vs. DR) | | | |
| g | %BE inside SWR | Two-way ANOVA with Bonferroni's multiple comparison test | sig | (CA1) group: F(1,1051)=1.435, P=0.231; session: F(1,1051)=3.910, P=0.048; group x session: F(1,1051)=6.476, P=0.011. Post-hoc: P=0.005 (Ctl: pre vs. post); P=0.628 (DR: pre vs. post); P=0.330 (pre: Ctl vs. DR); P=0.010 (post: Ctl vs. DR) | Ctl-pre: 15.096±1.171<br>Ctl-post: 19.773±1.459<br>DR-pre: 16.301±0.793<br>DR-post: 15.902±0.816 | Ctl-pre: 4.403±0.689<br>Ctl-post: 8.169±2.201<br>DR-pre: 3.298±0.421<br>DR-post: 8.429±1.623 | Ctl-pre: 10.951±1.258<br>Ctl-post: 23.798±4.377<br>DR-pre: 14.309±1.368<br>DR-post: 12.118±1.013 |
| | | | sig | (CA2) group: F(1,209)=0.108, P=0.742; session: F(1,209)=11.986, P=0.001; group x session: F(1,209)=0.282, P=0.596. Post-hoc: P=0.066 (Ctl: pre vs. post); P=0.001 (DR: pre vs.post) | | | |
| | | | sig | (CA3) group: F(1,438)=5.493, P=0.020; session: F(1,438)=9.007, P=0.003; group x session: F(1,438)=17.936, P<0.001. Post-hoc: P<0.001 (Ctl: pre vs. post); P=0.222 (DR: pre vs. post); P=0.123 (pre: Ctl vs. DR); P<0.001 (post: Ctl vs. DR) | | | |
